# Multi-layered regulation of neuroectoderm differentiation by retinoic acid in a primitive streak-like context

**DOI:** 10.1101/2021.03.18.435967

**Authors:** Luigi Russo, Hanna L. Sladitschek, Pierre A. Neveu

## Abstract

The formation of the primitive streak (PS) and the subsequent induction of neuroectoderm are hallmarks of gastrulation. Combining an *in vitro* reconstitution of this process based on mouse embryonic stem cells (mESCs) with a collection of knockouts in reporter mESC lines, we reassessed the contribution of retinoic acid (RA) signaling at early stages of neural commitment and its cross-talk with TGF*β* and Wnt signaling inhibition. Single-cell RNA sequencing analysis captured the temporal unfolding of cell type diversification from epiblast and primitive streak-like cells up to the emergence of anterior and posterior neural fates. In conditions thought to lack RA synthesis, we discovered a hitherto unidentified residual RA production via a sensitive RA reporter. Genetic perturbations proved that the RA-degrading enzyme Cyp26a1 safeguard the developmental capabilities of the PS-like cells, limiting neural differentiation in wild type and in *Chrd*^-/-^*Nog*^-/-^ or *Dkk1*^-/-^ cells. Finally, the knockout of the three RAR receptors highlighted their function as negative regulators of loci critical for neural induction. Overall, we identified two mechanisms whereby components of the RA pathway can control the formation of neural progenitors in our PS-like context: a RA-dependent neural induction gated by RARs, and a receptor-mediated repression in the absence of ligand.

## INTRODUCTION

During gastrulation, cells of the epiblast are allocated to the three germ layers (Tam and Behringer, 1997; Arnold and Robertson, 2009). Gastrulation is initiated by formation of the primitive streak (PS) and subsequent induction of neuroectoderm. Seminal experiments by Spemann and Mangold showed that the transplantation of the dorsal blastopore in amphibians could induce a secondary axis and neural tissue in the host embryo (Spemann and Mangold, 1924). This region or “organizer” secretes a range of molecules that mediate this induction (De Robertis, 2006). Among them, antagonists of the transforming growth factor b (TGFb) signaling pathway and in particular of bone morphogenetic proteins (BMPs) are considered pivotal for neuralization of the ectoderm (Hemmati-Brivanlou and Melton, 1997; Weinstein and Hemmati-Brivanlou, 1999). The inhibition of the Wnt signaling pathway is another potent inductive cue (Glinka et al., 1998). While most of the molecular mechanisms governing this process were determined in amphibians, they appear to be conserved in mammalian development (Levine and Brivanlou, 2007). Indeed, during mouse development the primitive streak and its distal tip, the node, possess organizer-like properties (Robb and Tam, 2004). The deletion of the two TGF*β* inhibitors Chordin and Noggin (Bachiller et al., 2000) or the knockout of the Wnt inhibitor Dkk1 (Mukhopadhyay et al., 2001) lead to the absence of anterior neural structures (forebrain) in mouse. Retinoic acid (RA) is another signaling molecule with potent neuralizing activity (Rhinn and Dollé, 2012) that was found to be produced by the Hensen’s node, the chick equivalent of the organizer (Hogan et al., 1992). Furthermore, RA signaling was detected in the primitive streak region of the mouse embryo at E7.5 (Rossant et al., 1991). At this developmental stage, Aldh1a2 is considered to be the only enzyme synthesizing RA from retinal (Rhinn and Dollé, 2012). Null mutations of *Aldh1a2*, in fact, lead to an absence of expression of an RA activity reporter (Niederreither et al., 1999), while the forebrain structures are still present. The described phenotype of the *Aldh1a2*^-/-^ embryos ruled out RA involvement in neural induction in early post-implantation development (Niederreither et al., 1999). This is in stark contrast with the widespread use of RA to induce neuronal fates from pluripotent cells *in vitro* (Ying et al., 2003; Bibel et al., 2004). Moreover, a role of RA in the formation of the posterior portion of the neural axis is well established. Here, the allocation of cell types to somite and spinal cord fates from bipotent neuromesodermal progenitors (NMPs) allows the extension of the body axis (Tzouanacou et al., 2009; Henrique et al., 2015; Kimelman, 2016). It was demonstrated that RA has a critical role in the differentiation of the bipotent progenitors to the neural lineage (Diez del Corral et al., 2003; Cunningham et al., 2016; Gouti et al., 2017). The effector of the developmental functions of RA is represented by the RAR family of nuclear receptors, which act as transcription factors regulated by RA (Dilworth and Chambon, 2001; Mic et al., 2003). RARs have the capability to either activate or repress the expression of target genes, recruiting co-regulators to the promoter regions containing their responsive elements (RAREs) (Perissi and Rosenfeld, 2005; Samarut and Rochette-Egly, 2012). While classical *in vivo* work established the importance of antagonizing TGF*β* and Wnt signaling pathways in the process of neural induction, and dismissed a contribution of RA signaling in this process, the molecular implementation of the neuroectoderm differentiation decision is largely unexplored. *In vitro* systems based on pluripotent stem cells enable to recapitulate crucial aspects of early post-implantation mammalian development (Shahbazi et al., 2019). Recent advances in the culture systems with micropatterned devices enabled the formation of germ layer fates in two-dimensional spatially ordered manner (Warmflash et al., 2014; Etoc et al., 2016). ESC aggregates allowed to differentiate in three dimensions demonstrated self-organizing properties, with formation of the three germ layers and axial elongation (van den Brink et al., 2014; Beccari et al., 2018; Simunovic et al., 2019; Moris et al., 2020), and the possibility to reach the somitogenesis stage in the presence of the appropriate extracellular matrix environment (van den Brink et al., 2020; Veenvliet et al., 2020).

Using an mESC-based system in which we can monitor the formation of neuroectoderm in the presence of a primitive streak-like population, we determined that neural induction by TGF*β* and Wnt antagonism could be impaired by inhibiting the RA receptors. Single-cell RNA sequencing captured the progression of the culture from epiblast and primitive streak-like cells to neural progenitors with different anteroposterior characteristics together with definitive endoderm and mesoderm cell types. By employing genetic and pharmacological interrogations in mESCs simultaneously reporting on different fates and the activation status of signaling pathways, we determined that the RA degrading enzyme Cyp26a1 limits the acquisition of neural fate, both in wild type cells and in the absence of the antagonists Chordin and Noggin or Dkk1. Using a highly sensitive RA reporter, we identified the presence of RA signaling in conditions commonly thought to lack RA synthesis ability. Finally, we identified a role of the RA receptors independent from RA in the control of the differentiation towards the neural lineage. Altogether, our results add valuable insights into the multi-layered regulation of RA signaling in the process of neuroectoderm formation.

## RESULTS

### Characterization of a system suitable for investigating the mechanisms of neuroectoderm formation

The primitive streak (PS) has organizer-like properties and the possibility to generate primitive streak-like cells *in vitro* should lead to the induction of a neuroectodermal fate among differentiation-competent cells (Figure 1A). To monitor the formation of PS-like cells and subsequent induction of neural progenitors, we used a double knock-in (2KI) reporter mESC line that we previously characterized (Sladitschek and Neveu, 2019) with *Sox1* locus targeted with GFP and *T* (also known as *Brachyury*) locus targeted with H2B-3xTagBFP. While *T* is a commonly used as a mesoderm marker, it is also expressed in the PS (Wilkinson et al., 1990). *Sox1* itself marks exclusively neural progenitors (Pevny et al., 1998). We previously showed that differentiation with the small molecule IDE1, which activates the TGF*β* pathway (Borowiak et al., 2009), determined the formation of a transient epiblast-like state and of intermediates that resembled the primitive streak (PS) in mouse embryos (Sladitschek and Neveu, 2019). Interestingly, we identified the presence in these cultures of putative neural progenitors (NPs) Sox1^GFP+^ cells along with T^TagBFP+^ cells, the candidate PS-like cells (Figure 1B).

**Figure 1.**
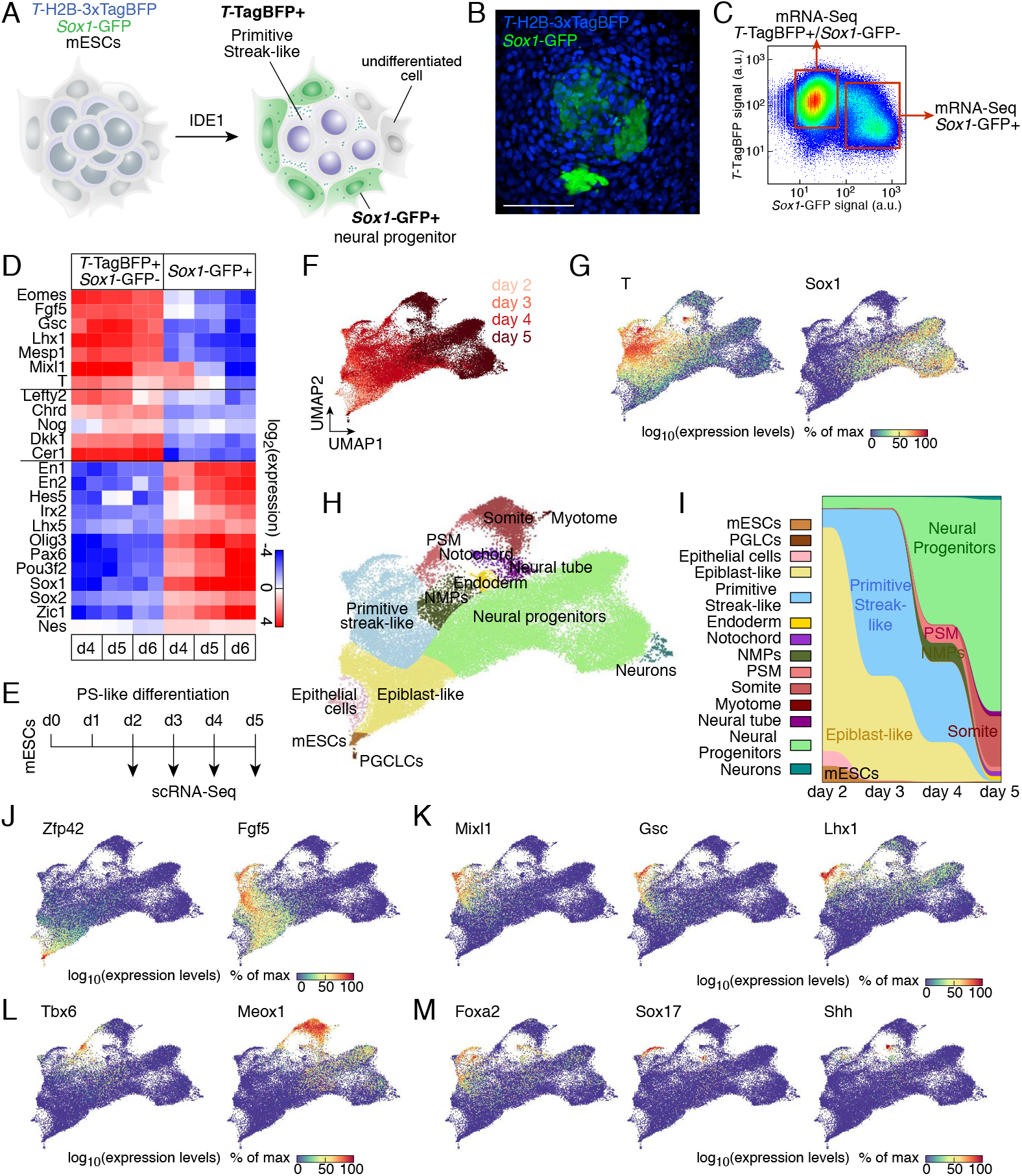
Characterization of neural induction by primitive streak-like cells at single cell level. (A) Scheme of the experimental principle to induce and monitor the formation of neuroectoderm by primitive streak-like cells using a double knockin mESC line reporting on *T* (also known as *Brachyury*) and *Sox1* expression. (B) *T*-TagBFP and *Sox1*-GFP reporter expression after 5 days of PS-like differentiation. Bar: 100 *μ*m. (C) Experimental strategy to characterize the different populations present in the differentiating culture. Populations are FACS-purified according to the reporter fluorescence and gene expression is measured by mRNA-Seq. (D) Expression levels (as measured by mRNA-Seq) of markers of the primitive streak or neural progenitors in FACS-purified populations after 4 to 6 days of PS-like differentiation. (E) Scheme of the experimental principle to temporally monitor PS-like differentiation using scRNA-Seq. (F) UMAP (uniform manifold approximation and projection) of 46,700 cells colored by the day of collection. (G) UMAP colored by the scaled expression of T and Sox1. (H) UMAP colored according to the identified populations (NMPs: Neuromesodermal progenitors, PSM: presomitic mesoderm, PGCLCs: primordial germ cell-like cells). (I) Alluvial plot representing the fraction of cells assigned to each fate indicated in (H) according to the day of differentiation. (J) UMAP colored by the scaled expression of the naïve pluripotency marker Zfp42 and the post-implantation epiblast marker Fgf5. (K) UMAP colored by the scaled expression of primitive streak markers. (L) UMAP colored by the scaled expression of presomitic mesoderm and somite markers. (M) UMAP colored by the scaled expression of endoderm and notochord markers. See also Figure S1.

We assessed the composition of the differentiating culture every day by flow cytometry. The increase in the TagBFP signal detected by the third day (Figure S1A) matched the increase in T mRNA levels as measured by mRNA-Seq (Figure S1B). In addition, we found that Sox1^GFP+^ cells could be detected in increasing numbers from day 3 (Figure S1C), mirroring the gradual increase in Sox1 mRNA levels (Figure S1D). Cell density played a role in the formation of the putative neural progenitors. In fact, initiating the differentiation with increasing cell numbers enhanced the fraction of Sox1^GFP+^ cells (Figure S1E).

To determine the identity of the different cell populations, we FACS-purified cells expressing TagBFP and GFP and measured their transcriptome by deep sequencing (Figure 1C). T^TagBFP+^ cells expressed markers associated with post-implantation epiblast and primitive streak fates such as Fgf5, T, Mixl1 and Goosecoid (Figure 1D). More importantly, the expression of the secreted antagonists that are commonly associated with the *in vivo* organizer was selectively higher in the T^TagBFP+^ population. Among these were the TGF*β* antagonists Chordin (Chrd) and Noggin (Nog) and the Wnt antagonist Dkk1. The Sox1^GFP+^ cells expressed a diverse range of neural progenitor markers, among which Sox2, Pax6 and Nestin (Figure 1D), confirming their neuroectodermal identity.

### Time resolved scRNA-Seq characterization of PS-like differentiation and neuroectoderm formation

A defining feature of our PS-like differentiation protocol was the coexistence of different fates, necessitating a finer characterization of the cellular heterogeneity than bulk mRNA-Seq. We therefore conducted single-cell RNA sequencing on differentiating cultures between day 2 and day 5 (Figure 1E). In total, we obtained expression profiles for 46,700 cells. Uniform manifold approximation and projection (UMAP) analysis showed only minimal overlap between consecutive days (Figures 1F and S1F). Profiles of FACS-purified T^TagBFP+^/Sox1^GFP-^ and Sox1^GFP+^ cells projected according to the respective expression territories of T and Sox1 (Figures 1G and S1G). Notably, cells with high T expression and Sox1-expressing cells formed distinct populations (Figure 1G). We could identify 35 cell subpopulations (Figures 1H and S1H) that were stratified by a number of markers (Figure S1I). Comparing the population distributions at each day, we observed a prevalence of epiblast and primitive streak fates at early time points (till day 3), followed by the formation of neural progenitors and PS derivatives later on (Figure 1I).

Naïve pluripotency markers were found in a subpopulation of day 2 cells while the other cells at this stage initiated the expression of primed epiblast markers (Figure 1J). Pluripotency factors had distinct behaviors: while Pou5f1 (also known as Oct4) expression was retained till day 4 in the epiblast and primitive streak lineages, Nanog expression was transiently reactivated in the PS-like population (Figure S1J). Sox2 was downregulated in PS-like cells and their derivatives, whereas its expression was maintained in NPs (Figure S1J). These findings faithfully recapitulated the expression patterns of these genes observed in E7.0 mouse embryos (Peng et al., 2019), and their proposed additional function as lineage priming factors (Thomson et al., 2011). PS markers were expressed in different subpopulations (Figures 1K and S1I), corresponding to different regions of the *in vivo* PS. Noteworthy, the expression of T encompassed both *bona fide* primitive streak and post-implantation epiblast cells, with transcripts level higher in the former (Figures 1G and S1I). Thus, the PS-like population marked by the expression of the TagBFP reporter at day 3 comprised a mixture of these two fates.

As the differentiation proceeded, the epiblast component was rapidly replaced by other fates and PS-derivatives started to be formed in our cultures (Figure 1I). Indeed, presomitic mesoderm and distinct subsequent somite fates could be found at days 4 and 5 (Figures 1L and S1I). Around the some time, a population resembling neuromesodermal progenitors (NMPs) also appeared (Figures 1I and S1I). Moreover, endoderm and notochord fates were present by day 5 (Figure 1M). Neuroectodermal cell types gradually accumulated in culture, at the expense of epiblast and primitive streak fates (Figures 1I and S1K). The existence of several NP populations (Figures S1H and S1I) was consistent with our bulk mRNA-Seq results (Figure 1D). Thus, our *in vitro* system recapitulated the fate diversification occurring in post-implantation embryos and notably the temporal evolution of the primitive streak *in vivo*.

### *In vitro* reconstitution of neural induction by Wnt and TGFβ antagonists

The inhibition of TGF*β* or Wnt signaling was shown to be critical for the induction of the neuroectodermal fate from pluripotent epiblast in the embryo (De Robertis, 2006). We sought to recapitulate the known action of these molecules in our *in vitro* model system by applying inhibitors to the differentiating cultures once the T^TagBFP+^ population was established at day 3 (Figure 2A). Adding a small molecule antagonist of the TGF*β* pathway SB431542 or blocking Wnt signaling using the tankyrase inhibitor XAV939 led to an increase in Sox1^GFP+^ cells (Figure 2B). Induction of Sox1^GFP+^ cells could be recapitulated as well by using recombinant versions of the secreted antagonists Noggin, DKK1 or secreted frizzled-related protein sFRP1(Figure S2A). Thus, inhibitors of either signaling pathway were potent inducers of neural fate in our *in vitro* system. We next sought to test whether the endogenous levels of the secreted antagonists were critical for the formation of the Sox1^GFP+^ cells. Therefore, we impaired the antagonists function generating *Chrd*^-/-^*Nog*^-/-^ mESCs and *Dkk1*^-/-^ mESCs in the 2KI background (Figures 2C). The PS-like differentiation of the knockout cells showed that the formation of Sox1^GFP+^ neural progenitors was severely compromised (Figure 2D). Enhancing TGF*β* or Wnt pathways activation by adding their agonists had similar overall effect of reducing the formation of neuroectodermal fate (Figures S2D and S2E), particularly evident when activating TGF*β* signaling. These results argue for a fine tuning of the formation of neuroectoderm dependent on the balance between the levels of agonist and inhibitors of TGF*β* and Wnt pathways.

**Figure 2.**
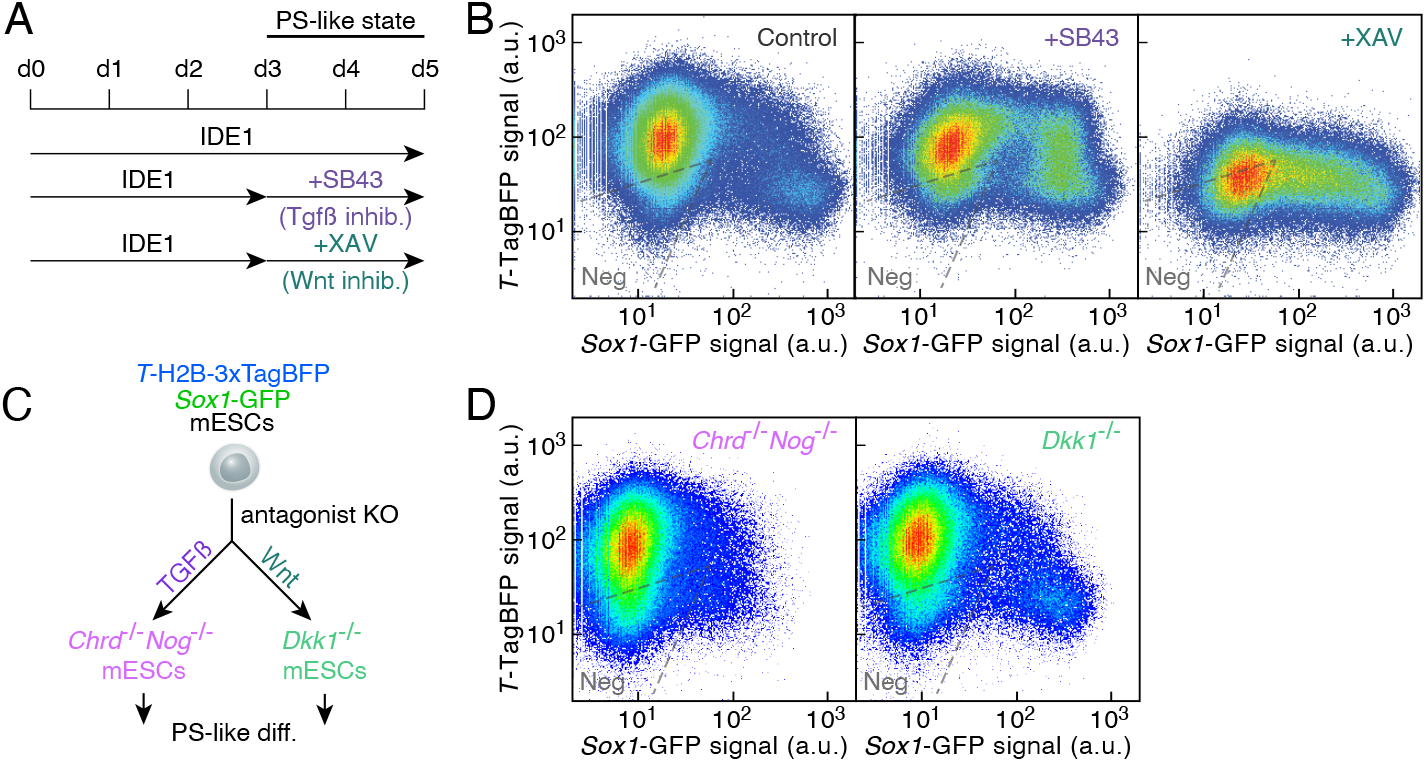
*In vitro* reconstitution of neural induction by Wnt and TGFβs antagonists. (A) Experimental strategy to assess the impact of TGF*β* or Wnt pathway inhibition on fate induction. SB431542 (SB43) inhibits TGF*β* receptors and XAV939 (XAV) is a tankyrase inhibitor. (B) *T*-TagBFP and Sox1-GFP reporter expression as measured by flow cytometry in control cultures (left panel) or in cultures treated with the TGF*β* inhibitor (SB43, middle panel) or the Wnt pathway inhibitor XAV (right panel). (C) Experimental strategy to assess the phenotype of cell lines with knockouts of antagonists of the TGF*β* and Wnt signaling pathways. (D) *T*-TagBFP and Sox1-GFP reporter expression after PS-like differentiation of *Chrd*^-/-^ *Nog*^-/-^ (left panel) or *Dkk1*^-/-^ (right panel) mESCs. See also Figure S2.

Our reporter line did not allow to resolve the exact fate that was promoted by the increased activation or the reduced inhibition of these pathways. We thus differentiated mESCs with Activin A or CHIR and compared the resulting expression profiles with the ones of T^TagBFP+^ cells induced by PS-like differentiation or with the ones of Epiblast Stem Cells (EpiSCs) whose maintenance relies on Activin A and FGF2 (Brons et al., 2007; Tesar et al., 2007) (Figure S2F). Principal component analysis revealed that TGF*β* and Wnt activation led to two orthogonal gene expression signatures (Figure S2G). During differentiation, the PS-like cells progressed from a state resembling the Activin A-treated cells towards a state in which Wnt signaling was more present (Figure S2G). Interestingly, the gene expression profiles of T^TagBFP+^ cells obtained from the PS-like differentiation of *Chrd*^-/-^ *Nog*^-/-^ mESCs or *Dkk1*^-/-^ mESCs projected towards the CHIR-differentiated samples (Figures S2H and S2I). For both cell lines, the Wnt signaling pathway was among the pathways with highest enrichment of differentially expressed genes as compared to wild type cells. The expression profiles of the *Chrd*^-/-^ *Nog*^-/-^ or *Dkk1*^-/-^ cells showed a downregulation of genes characteristic of PS and endoderm fates and an upregulation of presomitic mesoderm and somite markers as compared to the Activin A-differentiated cells (Figure S2J). Altogether, the absence of TGF*β* or Wnt antagonists led to effects comparable to the ones elicited by elevated Wnt signaling, favoring presomitic mesoderm and somite fates.

### Neural progenitors expressing a diverse range of markers emerge in the PS-like differentiation

The scRNA-Seq analysis evidenced an underlying heterogeneity within the population of neural progenitors (NPs) arising in the PS-like differentiation (Figure 3A). Indeed, different anteroposterior identities could be assigned to the NP subtypes according to the expression of markers characteristic of anterior neural tissues (*Hesx1*), anterior hindbrain (*Egr2* and *Hoxa2*), posterior hindbrain (*Hoxd4*), and spinal cord (*Hoxb9*) (Figures 3A) (Gouti et al., 2014). We hypothesized that the coexistence of multiple mechanisms of neural induction in our system could explain the expression of markers of neural progenitors with distinct developmental origin. To test this we compared the transcription profile of Sox1^GFP+^ cells derived from PS-like differentiation with the one of NPs generated differentiating mESCs in defined conditions such as Wnt signaling inhibition, TGF*β* inhibition and Wnt activation, or the well-established neural inducer retinoic acid (RA) (Ying et al., 2003) (Figure 3B). The NPs induced by each differentiation regime expressed distinct sets of transcription factors that spanned the set upregulated in the heterogenous PS-induced NP population (Figure 3C). Wnt inhibition led to the upregulation of anterior neuroectodermal markers such as Lhx5, Otx2 and Six3 (Figure 3C). Increased expression of markers of the posterior portion of the neural axis was found upon TGF*β* inhibition and Wnt activation (Figure 3C). Nevertheless, these two different NP-induction methods did not account for the full complexity of the expression profile of NPs obtained in the PS-like differentiation. Indeed, upregulation of markers such as Brn1, Brn2 or Irx3, could only be recapitulated by RA treatment (Figure 3C). This led us to infer that RA signaling might be in part responsible for the formation of neural progenitors in our heterogeneous system.

**Figure 3.**
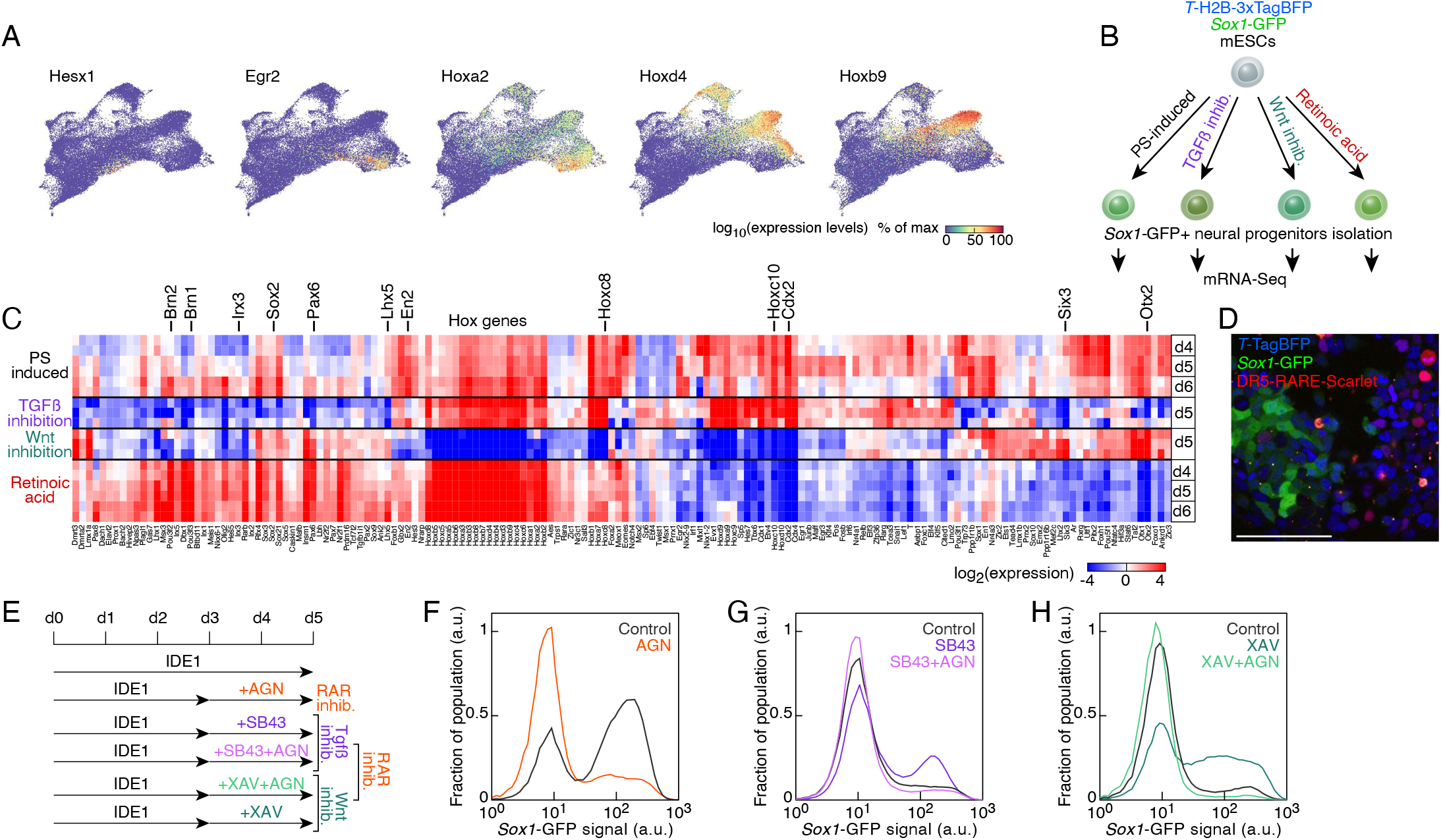
Retinoic acid RA signaling underlies the formation of neural progenitors in the PS-like differentiation. (A) UMAP colored by the scaled expression of neuroectodermal markers, ordered by their expression along the anteroposterior axis *in vivo*. (B) Scheme of the experimental principle to characterize neural progenitors induced by PS-like cells, TGF*β* or Wnt pathway inhibition or retinoic acid (RA) treatment. (C) Expression levels of transcription factors and regulators differentially expressed in Sox1^GFP+^ cells. (D) Reporter expression after 5 days of differentiation of a double knockin *T*-TagBFP and *Sox1*-GFP mESC line transgenic for a DR5-based retinoic acid signaling reporter. Bar: 100 *μ*m. (E) Experimental strategy to assess the crosstalk between RA signaling and TGF*β* or Wnt pathway inhibition on fate induction. AGN193109 (AGN) is a retinoic acid receptor (RAR) antagonist. (F) Sox1-GFP expression levels after PS-like induction with RA signaling inhibition (black: control, orange: AGN). (G) Sox1-GFP expression levels after PS-like induction with TGF*β* and RA signaling inhibition (black: control, purple: SB43, pink: SB43 and AGN). (H) *Sox1*-GFP expression levels after PS-like induction with Wnt and RA signaling inhibition (black: control, teal blue: XAV, light green: XAV and AGN). See also Figure S3.

### RA signaling underlies the formation of neural progenitors in the PS-like differentiation

As a proof of concept of the RA ability to induce neural fate in our differentiation system, we added RA to the medium at pharmacological concentrations after the PS-like population was established (Figures S3A). While most of the cells proceeded with acquiring the PS program in standard differentiation condition, the RA completely diverted the fate decision towards neuroectoderm formation, as expected (Figures S3B). The differentiation regime with the small molecule IDE1 contains serum at low concentrations. RA precursors present in the serum could be converted by the cells in the final product RA. To test the presence of RA signaling, we stably inserted in the 2KI line a fluorescent reporter construct relying on the well-established DR5-based RA response element (RARE) (Rossant et al., 1991) controlling the expression of the fluorescent protein Scarlet. Scarlet positive cells could be detected from day 3 of the PS-like differentiation onwards (Figure 3D and S3C), demonstrating the presence of retinoic acid signaling. This paralleled the identification of RA signaling in the primitive streak region of wild type E7.5 mouse embryos through a reporter relying on the same RARE (Rossant et al., 1991).

We went on testing the effects of perturbing RA signaling on the formation of neural progenitors. Adding to the differentiation medium increasing amounts of the RA precursor vitamin A (also known as retinol) enhanced the formation of Sox1^GFP+^ cells (Figure S3D). This was accompanied by an increase in RA signaling, as captured by our RA signaling reporter line (Figure S3E). Conversely, pharmacological inhibition of the RA receptors (RARs) with the small molecule AGN193109 completely prevented Scarlet expression (Figure S3F) and decreased the fraction of Sox1^GFP+^ cells (AGN, Figure 3E and Figure 3F). The extent of the reduction of neural progenitors could be enhanced by initiating RAR inhibition at early differentiation time points (Figure S3G). The observation implied that the inhibition of the RA receptors prevented the formation of neuroectodermal cells, but did not hamper the neural progenitors already present in the culture. The extent of this effect prompted us to test the existence of cross-talk between RA signaling and the mechanism of neural induction by TGF*β* or Wnt inhibition. We therefore applied simultaneously the RAR antagonist AGN and TGF*β* or Wnt inhibitors (Figure 3E). Surprisingly, AGN prevented the increase of the Sox1^GFP+^ population normally associated with the inhibition of either of the two pathways (Figures 3G and 3H). This result indicated that in the cascade of events leading to neuroectoderm formation, the RA receptors could control a step downstream of TGF*β* or Wnt inhibition.

### Aldh1a2-independent RA signaling

We found that components of the RA signaling enabled the induction of neuroectoderm elicited by the antagonists of the TGF*β* or Wnt pathways. Only the RA precursor, vitamin A, was present in our differentiation medium, therefore, the final product had to be synthesized by the cells themselves. The oxidation of retinal in retinoic acid is performed by the retinaldehyde dehydrogenases Aldh1a1, Aldh1a2 or Aldh1a3 (Rhinn and Dollé, 2012). According to our mRNA-Seq data, Aldh1a2 was upregulated during PS-like differentiation, whereas Aldh1a1 and Aldh1a3 expressions did not exceed background levels (Figure S4A). Aldh1a2 transcripts levels were particularly elevated in T^TagBFP+^ cells compared to Sox1^GFP+^ cells (Figure S4B). This mirrored the known pattern of expression of these three genes in post-implantation mouse embryos, particularly the expression of Aldh1a2 in the primitive streak at E7.5 (Ribes et al., 2009).

A contribution of RA signaling in the generation of anterior neural structures was excluded by the phenotype of *Aldh1a2*^-/-^ mouse embryos, exhibiting forebrain structures in the absence of detectable expression of a DR5-based RA reporter at E7.5 (Niederreither et al., 1999). This result based on mouse genetics seemed at first glance in contradiction with our finding that the induction of neuroectoderm by TGF*β* and Wnt inhibition needed active RA signaling. Therefore, we sought to reproduce *in vitro* the *Aldh1a2*^-/-^ condition and to investigate whether RA signaling was completely abolished by *Aldh1a2* ablation. We generated *Aldh1a2*^-/-^ mESCs (Figure S4C) in the 2KI line bearing the RA activity reporter relying on the same DR5-RARE used in the mouse model. PS-like differentiation of *Aldh1a2^-/-^* mESCs led to the formation of Sox1^GFP+^ cells despite the absence of DR5-RARE-Scarlet+ cells in the culture (Figure 4A and S4D). This result was in accordance with the phenotype of *Aldh1a2*^-/-^ mouse embryos (Niederreither et al., 1999). In order to assess whether the *Aldh1a2*^-/-^ cells were completely devoid of RA production, we provided extra precursor for its synthesis, vitamin A, or inhibited RARs during PS-like differentiation (Figure 4B). In the absence of RA synthesis these treatments should elicit no effect, that is the fraction of Sox1^GFP+^ cells should be insensitive to perturbation to RA signaling. Nevertheless, the RAR antagonist AGN led to a decrease of the fraction of Sox1^GFP+^ population (Figure 4C). Surprisingly, upon addition of vitamin A the majority of the cells expressed the GFP reporter (Figure 4C) and the DR5-RARE-Scarlet reporter could now be detected in the *Aldh1a2*^-/-^ differentiating cultures (Figure S4D). Altogether, the results demonstrated that the genetic ablation of Aldh1a2 was not sufficient to fully abrogate RA signaling. The DR5-RARE reporter used in previous studies was unable to capture the residual RA levels in *Aldh1a2*^-/-^ cultures.

**Figure 4.**
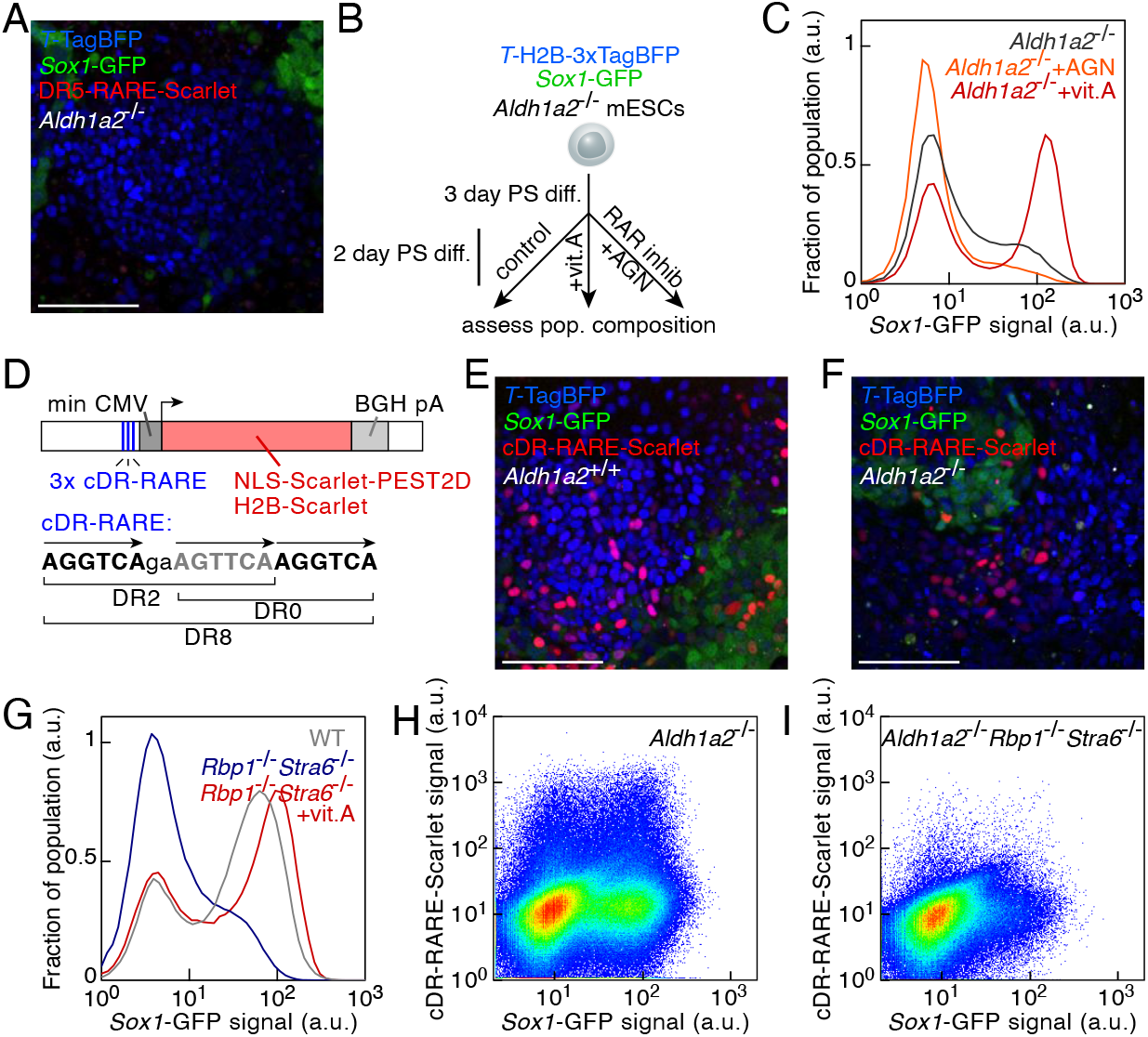
Aldh1a2-independent RA signaling during PS-like differentiation. (A) Reporter expression after 5 days of PS-like differentiation of a double knockin *T*-TagBFP *Sox1*-GFP *Aldh1a2*^-/-^ mESC line transgenic for a RA signaling reporter (DR5-RARE-Scarlet). Bar: 100 *μ*m. (B) Scheme of the experimental principle to monitor the impact of perturbing RA signaling on the PS-like differentiation of *Aldh1a2*^-/-^ mESCs. (C) Sox1-GFP reporter expression in *Aldh1a2*^-/-^ cultures after PS-like differentiation (black: control, orange: AGN from day 3, red: vitamin A from day 3). (D) Scheme of an RA-responsive transcriptional reporter relying on three RA responsive elements (RARE) consisting of three RAR binding sites creating direct repeats (DR) with three different spacing combinations (cDR: composite direct repeat, min CMV: minimal CMV promoter, BGH pA: bovine growth hormone poly A). (E) Reporter expression after 5 days of PS-like differentiation of a double knockin *T*-TagBFP Sox1-GFP wild type (*Aldh1a2*^+/+^) mESC line transgenic for the cDR RA signaling reporter (cDR-RARE-Scarlet). Bar: 100 *μ*m. (F) Reporter expression after 5 days of PS-like differentiation of a double knockin *T*-TagBFP Sox1-GFP *Aldh1a2*^-/-^ mESC line transgenic for the cDR RA signaling reporter (cDR-RARE-Scarlet). Bar: 100 *μ*m. (G) Sox1-GFP reporter expression after PS-like differentiation of wild type (gray) or *Rbp1*^-/-^ *Stra6*^-/-^ cells without (blue) or with (red) additional vitamin A. (H, I) Sox1-GFP and cDR-RARE-Scarlet reporter expression after PS-like differentiation of *Aldh1a2*^-/-^ (H) or *Aldh1a2*^-/-^ *Rbp1*^-/-^ *Stra6*^-/-^ (I) mESCs. See also Figure S4.

We wondered whether *Aldh1a2*^-/-^ cells were still able to respond to the RA levels produced by wild type cultures and, vice versa, whether *Aldh1a2*^+/+^ cells’ response to RA would be affected by the presence in culture of cells unable to synthesize RA. To address both questions we set up a co-culture experiment by mixing wild type *Aldh1a2*^+/+^ cells and mutant *Aldh1a2*^-/-^ cells and differentiating them together (Figure S4E). The two genotypes could be distinguished by the constitutive expression of an additional fluorescent protein H2B-2xiRFP670. Under such conditions, DR5-RARE-Scarlet+ cells were found in the *Aldh1a2*^-/-^ fraction at a rate comparable to the one in *Aldh1a2*^+/+^ cells (Figures S4F and S4G). This proved that RA signaling occurred in a paracrine fashion in our *in vitro* system. On the other hand, the fraction of *Aldh1a2*^+/+^ cells expressing the RA reporter was reduced in the co-culture setting (Figure S4G) as compared to pure wild type culture (Figure S4F). This observation implied that a cell’s response to RA did not depend on its own RA production but rather on the overall level present in the media.

### A highly sensitive RA reporter captures Aldh1a2-independent RA signaling

Our results stressed that the traditional DR5-based RARE might capture only a subset of conditions in which RA signaling was present. A composite RARE was found to have much higher affinity for RARs than the traditional DR5 binding site (Moutier et al., 2012). This composite RARE consisted of three RAR binding sites forming direct repeats with three different spacings (Figure 4D). We developed a fluorescent reporter construct relying on this RARE (which we termed cDR-RARE) driving the expression of the fluorescent protein Scarlet and we stably inserted it in the 2KI line (Figure 4D). The reporter could detect sub-nanomolar concentrations of exogenously applied RA (Figure S4H). Many more cDR-RARE-Scarlet+ cells could be detected during PS-like differentiation of the new reporter line (Figure 4E) compared to the traditional DR5-based reporter (Figures S4I and S4J). As expected, Scarlet expression could be blocked by the RAR antagonist AGN (Figure S4K). Crucially, this reporter was able to capture the presence of RA signaling in *Aldh1a2*^-/-^ cells (Figures 4F and S4L). Therefore, a fraction of RA was produced in an Aldh1a2-independent manner at sufficient levels to be detected by the cDR-RARE reporter and to impact on the formation of Sox1^GFP+^ cells.

### Vitamin A availability regulates RA signaling levels during PS-like differentiation

Providing increasing amounts of vitamin A led to increasing numbers of Sox1^GFP+^ cells (Figure S3D), indicating that the cells were sensitive to the external vitamin A concentration. Interestingly, the transcript levels of the cellular retinol binding protein Rbp1 and the Rbp-receptor Stra6, a major mediator of the cellular uptake of vitamin A (Kawaguchi et al., 2007), were upregulated during PS-like differentiation (Figures S4M and S4N). Rbp1 binds to retinol and it is thought to increase its concentration within the cell protecting it from rapid degradation and helping RA synthesis (Ghyselinck et al., 1999; Napoli, 2016). Elevated expression levels of Rbp1 were found in the primitive streak of E7.0 mouse embryos as determined by spatial transcriptomics (Peng et al., 2019). We hypothesized that vitamin A uptake through Stra6 and intracellular storage by Rbp1 could be an important parameter in controlling RA levels. Therefore, we generated double knockout *Rbp1*^-/-^ *Stra6*^-/-^ mESCs (Figure S4O) and submitted them to PS-like differentiation. These cells were impaired in generating Sox1^GFP+^ cells compared to wild type cells (Figure 4G). However, the defect could be reversed by increasing vitamin A concentration in the medium (Figure 4G). We next asked whether disrupting the cellular supply of vitamin A would further decrease the RA signaling levels in *Aldh1a2*^-/-^ cells that were impaired in RA synthesis. PS-like differentiation of the triple knockout *Aldh1a2*^-/-^ *Rbp1*^-/-^ *Stra6*^-/-^ cells displayed indeed reduced RA signaling as measured by the cDR-RARE-based fluorescent reporter and a decreased fraction of Sox1^GFP+^ cells compared to *Aldh1a2*^-/-^ cells (Figures 4H and 4I). This demonstrated that the control of the intracellular vitamin A levels by the Stra6-Rbp1 axis contributed to determine the eventual levels of RA signaling.

### Cyp26a1 limits RA signaling during PS-like differentiation

We next sought to test whether the regulation of RA degradation helped define the subset of cells that can be responsive to RA within the differentiation competent PS-like population. RA oxidation to inactive metabolites is mediated by cytochrome P450 Cyp26 enzymes (Rhinn and Dollé, 2012). Cyp26a1 mRNA levels were highly upregulated during PS-like differentiation (Figure S5A), and particularly in the T^TagBFP+^ cells as compared to Sox1^GFP+^ cells (Figure S5B). Cyp26a1 upregulation in our culture system mirrored its expression in the primitive streak at E7.0 *in vivo* (Fujii et al., 1997). We hypothesized that the RA degrading enzyme Cyp26a1 could hamper neural induction by limiting RA levels. Therefore, we introduced null mutations in the *Cyp26a1* locus in the 2KI-RA reporter line, which provided a convenient read out of the effects of this perturbation on RA signaling and neuroectoderm formation (Figure S5C). As expected, *Cyp26a1*^-/-^ cells formed more Sox1^GFP+^ cells and displayed elevated RA signaling as measured with the RA activity reporter (Figure 5A). The loss of Cyp26a1 further enhanced the formation of neuroectodermal cell types at the expense of other fates in response to extracellular vitamin A (Figure S5D).

**Figure 5.**
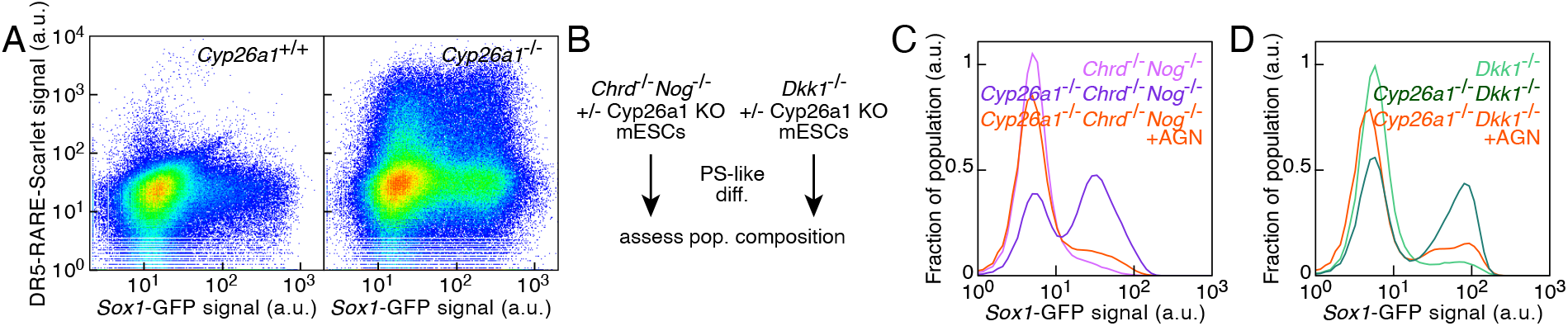
Cyp26a1 is a key factor limiting RA levels and neuroectoderm differentiation during PS-like differentiation. (A) *Sox1*-GFP and DR5-RARE-Scarlet reporter expression after PS-like differentiation of wild type (left panel) or *Cyp26a1*^-/-^ cells (right panel). (B) Scheme to assess the interplay between Cyp26a1-mediated dampening of RA signaling and TGF*β* or Wnt signaling. (C) Sox1-GFP reporter expression in *Chrd*^-/-^ *Nog*^-/-^ (pink) or *Cyp26a1*^-/-^ *Chrd*^-/-^ *Nog*^-/-^ (purple, orange: AGN added after day 3) cultures after PS-like differentiation. (D) Sox1-GFP reporter expression in *Dkk1*^-/-^ (light green) or *Cyp26a1*^-/-^ *Dkk1*^-/-^(dark green, orange: AGN added after day 3) cultures after PS-like differentiation. See also Figure S5.

We wondered whether Cyp26a1-mediated RA degradation affected the capability of the cells to commit to neural fate in response to different RA levels (Figure S5E). Differentiating wild type and *Cyp26a1*^-/-^ cells in chemically defined medium with low (1nM) RA concentration, evidenced a large difference in the fraction Sox1^GFP+^ cells (Figure S5F). The majority of the cells devoid of the RA degrading enzyme, indeed, acquired the neural fate in contrast to a minor fraction of *Cyp26a1*^+/+^ (Figure S5F). High RA concentrations could instead overcome the threshold imposed by Cyp26a1, so that *Cyp26a1*^-/-^ and wild type cells displayed similar capacity to differentiate to neuroectoderm in this condition (Figure S5G). The absence of Cyp26a1, thus, increased the response to low concentrations of RA. Altogether, Cyp26a1 played a key role in reducing RA levels and the acquisition of neural fate during PS-like differentiation.

### RA degradation by Cyp26a1 limits neuroectoderm formation in the *Chrd*^-/-^ *Nog*^-/-^ and *Dkk1*^-/-^ cells

We showed that removing the RA-degrading enzyme Cyp26a1 increased the response to RA signaling and the fraction of neuroectodermal cells generated during the PS-like differentiation. On the contrary, null mutations of *Chrd* and *Nog* or *Dkk1* impaired the formation of neural progenitors. We sought to test whether the reduced neuroectoderm formation due to the absence of the TGF*β* or Wnt inhibitors could be counteracted by the loss of Cyp26a1. Therefore, we generated mESCs lacking Cyp26a1 and either TGFb or Wnt antagonists and subjected them to PS-like differentiation (Figures 5B, S5H and S5I). Triple knockout *Chrd*^-/-^ *Nog*^-/-^ *Cyp26a1*^-/-^ and double knockout *Dkk1*^-/-^ *Cyp26a1*^-/-^ cells originated a large fraction of Sox1^GFP+^ cells compared to *Chrd*^-/-^ *Nog*^-/-^ cells or *Dkk1*^-/-^ cells (Figures 5C, 5D and S5J). This increase could be reverted by the RAR antagonist AGN, underlining that this effect was strictly dependent on RA signaling (Figures 5C and 5D). Therefore, Cyp26a1-mediated dampening of RA signaling proved critical in reducing the exposure of the PS-like population to the differentiating effects of RA they produce.

### RA signaling status accounts for neural progenitor diversity

We showed conditions in which RA signaling was impaired (AGN-treated cells and *Aldh1a2*^-/-^ cells) and others where RA signaling was active (cells expressing the cDR-RARE-Scarlet reporter). Thus, we sought to characterize the gene expression changes associated with active or inactive RA signaling both in T^TagBFP+^ and Sox1^GFP+^ subpopulations (Figure 6A). Expression changes observed in the T^TagBFP+^ cells upon AGN treatment were well correlated with the ones observed in *Aldh1a2*^-/-^ cells (Figure 6B) and were anticorrelated with the expression changes due to active RA signaling in the cDR-RARE-Scarlet^+^ T^TagBFP+^ cells (Figure 6C). Active RA signaling could either repress (Figure S6A) or increase (Figure S6B) gene expression. Interestingly, there was no correlation between the changes in gene expression associated with RA signaling and the ones between Sox1^GFP+^ and T^TagBFP+^ cells (Figure S6C). This suggested that RA was involved in early steps of neural commitment and not responsible for the final expression signature of Sox1^GFP+^ cells.

**Figure 6.**
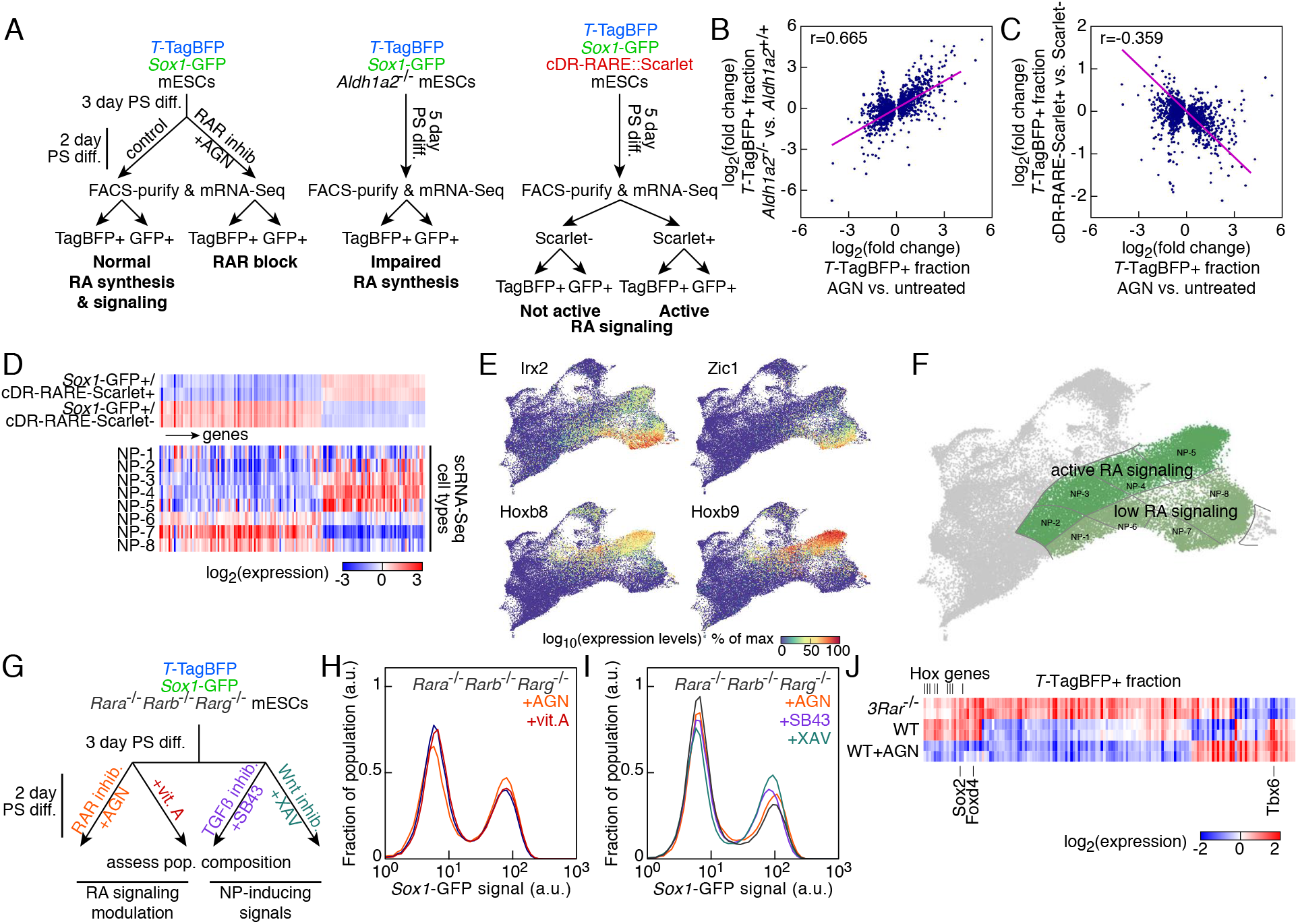
Role of RA signaling and RAR receptors in neural commitment and in establishing neural progenitor diversity. (A) Experimental strategy to assess the impact of RA signaling on gene expression during PS-like differentiation both in the T^TagBFP+^ (TagBFP+) and the Sox1^GFP+^ (GFP+) populations. (B) Comparison of differential expression in T^TagBFP+^ cells after AGN treatment or in *Aldh1a2*^-/-^ cells (Pearson’s r=0.665, p=10^-181^). (C) Comparison of differential expression in T^TagBFP+^ cells after AGN treatment or in cDR-RARE-Scarlet+ cells with active RA signaling (Pearson’s r=-0.359, p=2.10^-44^). (D) Expression levels of genes with differential expression between cDR-RARE-Scarlet+ and Scarlet-subpopulations of the Sox1^GFP+^ fraction after PS-like differentiation and in the 8 neural progenitor (NP) categories identified by scRNA-Seq. (E) UMAP colored by the scaled expression of markers identifying different NP territories. (F) UMAP with the RA signaling status highlighted in the NP territories. (G) Experimental strategy to monitor the impact of RA signaling modulation or NP-inducing cues in *Rara*^-/-^ *Rarb*^-/-^ *Rarg*^-/-^ mESCs. (H) Sox1-GFP reporter expression in *Rara*^-/-^ *Rarb*^-/-^ *Rarg*^-/-^ cultures after PS-like differentiation with perturbation of RA signaling (black: control, orange: AGN from day 3, red: vitamin A from day 3). (I) Sox1-GFP reporter expression in *Rara*^-/-^ *Rarb*^-/-^ *Rarg*^-/-^ cultures after PS-like differentiation with TGF*β* or Wnt signaling inhibition (black: control, orange: AGN, purple: SB43, teal blue: XAV). (J) Expression levels of transcriptional factors with differential expression in the T^TagBFP+^ fraction after PS-like differentiation of *Rara*^-/-^ *Rarb*^-/-^ *Rarg*^-/-^ (*3Rar*^-/-^) cells or wild type cells without (WT) or with (WT+AGN) treatment with the RAR antagonist AGN. See also Figure S6.

We next identified genes differentially expressed between the Sox1^GFP+^ cells expressing the RA reporter and the Sox1^GFP+^ cells in which cDR-RARE-Scarlet was not detected. We assessed the expression of these genes in the eight different neural progenitor (NP) subpopulations identified by scRNA-Seq (Figures S1H and 6D). NP-2, NP-3, NP-4 and NP-5 resembled the Sox1^GFP+^ cells with active RA signaling, whereas the NP-6, NP-7 and NP-8 classes shared a signature with cDR-RARE-Scarlet negative cells (Figure 6D). Among the transcription factors differentially expressed were NP markers associated with distinct anteroposterior identity such as Irx2, Zic1 and Hoxb6 (Figure 6E). The NP-1 subpopulation presented elevated expression levels of genes characteristic of anterior NP identity (Figure S6D and S6E). These markers were upregulated upon AGN treatment as compared with the cells in which the RA signaling reporter was not detected (Figure S6F). Altogether, differences in RA signaling activation status could account for the particular signatures of the NP subpopulations identified by scRNA-Seq, broadly dividing them into two regions (Figure 6F).

### RAR knockout cells exhibit increased propensity to neuroectoderm differentiation

The RA receptors (RARs) are the transcriptional effectors of RA signaling (Chambon, 1996). To obtain a condition where RA signaling cannot be transduced, we derived mESCs lacking all three RARs (RAR-null cells) (Figure S6G). As expected and contrary to wild type cells, RA failed to induce the expression of the DR5-RARE-based reporter in *Rara*^-/-^ *Rarb*^-/-^ *Rarg*^-/-^ mESCs (Figures S6H and S6I). Despite the absence of RA signaling, RAR-null cells displayed the formation of a Sox1^GFP+^ population after PS-like differentiation. Therefore, we tested whether the induction of neural fate in these cells was responsive to alterations of RA signaling or to TGF*β* and Wnt inhibition (Figure 6G). The fraction of Sox1^GFP+^ cells appeared completely insensitive to the addition of vitamin A or a RAR antagonist (Figure 6H), differently from what observed for the *Aldh1a2*^-/-^ condition. The repression of TGF*β* or Wnt signaling displayed residual capacity to increase the fraction of neural progenitors in RAR-null cells (Figure 6I). The results underpin a clear functional distinction with regard to neuroectoderm formation between inhibiting the RA receptors with AGN and completely removing them in the RAR knockout.

We profiled the gene expression of the T^TagBFP+^ and Sox1^GFP+^ subpopulations of *Rara*^-/-^ *Rarb*^-/-^ *Rarg*^-/-^ cells and compared them with the ones of wild type cells or of cells treated with the RAR antagonist AGN. Interestingly, AGN-treated T^TagBFP+^ cells and T^TagBFP+^ RAR-null cells presented differences in transcription factor expression that might explain the increased propensity of the RAR-null cells towards the formation of Sox1^GFP+^ cells (Figure 6J). In fact, T^TagBFP+^ RAR-null cells exhibited reduced expression of Tbx6, a gene whose loss leads to the formation of neural tissue at the expense of somites *in vivo* (Chapman and Papaioannou, 1998). Moreover, these cells presented an upregulation of Sox2, whose misexpression in paraxial mesoderm was reported to cause ectopic neural tube formation (Takemoto et al., 2011). The Sox1^GFP+^ population originating from the RAR-null cells possessed as well a distinct expression profile (Figure S6J). Hox genes and members of the Cdx family were downregulated in these cells, whereas a number of transcription factors characteristic of neural fate (such as Neurog2, Neurod4 or Nhlh1) had elevated expression. Thus, the lack of RARs had distinct effects compared to their pharmacological inhibition and could not be assimilated to an absence of RA signaling. We inferred that the repressive function of the receptors in absence of their ligand could be a distinction of critical importance in the regulation of neuroectoderm formation. We tested this hypothesis, comparing the outcome of differentiating wild type and RAR-null cells in chemically defined medium devoid of RA precursors. The differentiation in absence of the RA receptors resulted in the formation of many more Sox1^GFP+^ cells than in the wild type condition, demonstrating the importance of RARs in the homeostasis of neuroectoderm formation (Figure S6K).

## DISCUSSION

In this work, we combined a system reproducing the maturation of primitive streak-like cells and the formation of both anterior and posterior neuroectodermal fates with a large collection of reporter mESC lines harboring genetic ablations of key signaling factors. In such a context, we identified the presence of crosstalk between RA signaling and the default mechanisms of neural induction by Wnt or TGF*β* inhibition, and we propose a more widespread contribution of multiple components of the RA pathway in the decision to undergo neuroectoderm differentiation.

While cell type heterogeneity is normally an obstacle for the applications of simple bidimensional culture systems, we exploited mESC capability to give rise to pletora of fates in response to a starting homogenous signaling environment. Our scRNA-Seq analysis, in fact, demonstrated the coexistence of a wide array of cell types in our culture despite the absence of defined geometrical constraints. Neither long-range order nor a three-dimensional arrangement were necessary to direct neighbouring cells towards different fates. Instead, fate diversity was created by local interactions between cells that generated signaling microenvironments. Methods ranging from precise patterning of bidimensional cultures to novel tridimensional cultures of ESC aggregates are proving successful in reproducing the self-organized spatially ordered fate diversification proper of embryonic development (Warmflash et al., 2014; van den Brink et al., 2020; Veenvliet et al., 2020). While our approach lacked the layer of complexity derived from geometrical confinement, the uncoupling between fate specification and tissue-scale patterning/morphogenesis allowed to address questions such as how the local control of the signaling environment can lead to the onset of different fates.

The scRNA-Seq analysis indicated that epiblast and primitive streak cell types prevailed in our culture at early time points (till day 3 of differentiation). Both populations were embedded into the T^TagBFP+^ PS-like cells, with higher level of expression of T in the *bona fide* primitive streak. During early stage of gastrulation, the formation of neural lineage begins from the epiblast cells not included in the primitive streak. This was the rationale behind starting most of the pharmacological perturbations of neural induction from day 3. The emergence of the streak *in vivo* marks also the beginning of anteroposterior patterning. Instances of this process could be found in our culture which did not allow axis formation. In fact, the PS-like cells gave rise to anterior/endoderm derivatives and gradually more posterior/mesoderm fates. Moreover, we identified the presence of neuromesodermal progenitors (NMPs), which fuel axial elongation *in vivo* and are precursors of both somites and spinal cord (Tzouanacou et al., 2009; Henrique et al., 2015; Kimelman, 2016). The diversification of the fates originating from the PS-like cells and the disappearance of epiblast cell types were accompanied by an increasing fraction of cell types allocated to the neural lineage. Noteworthy, the neural progenitors arising in culture spanned the entire neural axis as demonstrated by the expression of genes marking distinct anteroposterior territories. The importance of this finding lies in that it argues for multiple origins of the subpopulations of neural progenitors and multiple mechanisms of induction.

Neural induction by inhibition of Wnt or TGF*β* pathways occurs in the anterior epiblast of the mouse conceptus and was reproduced in our system. This was confirmed independently by the pharmacological inhibition of the pathways, and by impairing the endogenous levels of the antagonists. Indeed, Wnt inhibition led to the formation of neural progenitors expressing anterior markers, consistent with previous reports (Watanabe et al., 2005), whereas neuroectoderm with more posterior identity could be derived by inhibiting TGFb signaling while activating Wnt. The impaired neuroectoderm formation in *Chrd*^-/-^ *Nog*^-/-^ or *Dkk1*^-/-^ cells was in line with the corresponding mutant mouse embryos lacking anterior neural structures (Bachiller et al., 2000; Mukhopadhyay et al., 2001). We noticed that the reduction in the neural progenitors formed in the knockout cultures was accompanied by increase in Wnt signaling and formation of mesodermal cells. This was coherent with Wnt contribution to the formation of the mesoderm germ layer *in vivo* (Liu et al., 1999; Huelsken et al., 2000).

The inhibition of Wnt or TGF*β* signaling, however, could not reproduce the entire spectrum of anterior and posterior neuroectoderm obtained during the PS-like differentiation. In fact, a component of neural progenitors relied on RA signaling. We detected retinoic acid (RA) signaling activity in the PS-like differentiation generating a RA sensor line based on the same RA responsive elements (RAREs) used in the classical mouse model. The various NP subpopulations differed in their RA signaling status, with markers corresponding to anterior fates being expressed in cells with low RA signaling. This finding is in accordance with the proposed caudalizing effects of RA during development (Durston et al., 1989). More importantly, the mechanisms of neural induction at work in the PS-like culture were not independent from each other, one originating anterior NPs and the other posterior. Instead, the formation of neural progenitors elicited by pharmacological inhibition of Wnt or TGF*β* signaling could be hindered by blocking the RA receptors (RARs) with AGN. These observations suggested that the RA receptors might control a step downstream of Wnt and TGF*β* inhibition in the cascade of events leading to the acquisition of neural fate,.

The identification of all RA signaling with the DR5-RARE based reporter, together with the absence of expression of this reporter in the *Aldh1a2*^-/-^ embryos and their phenotype, played a central role in the definition of the functions of RA in neural specification (Niederreither et al., 1999). The mRNA levels of the retinaldehyde dehydrogenase Aldh1a2 were strongly upregulated during PS-like differentiation and Aldh1a2-mediated synthesis was crucial to generate high levels of RA signaling. However, Aldh1a2 itself did not account for all the RA production. The ability to perturb RA signaling in *Aldh1a2*^-/-^ cells enabled us to discover residual RA production that could not be captured by a classical RA reporter. In the knockout cells, in fact, RA signaling could be detected again and neuroectoderm formation could be enhanced by increasing the amount of substrate for RA synthesis (vitamin A). Developing a highly sensitive RA activity reporter, we demonstrated the presence of RA signaling during PS-like differentiation of *Aldh1a2*^-/-^ cells. While this result awaits confirmation in future *in vivo* studies, its significance lies in the identification of alternative ways to respond to RA beyond the *Aldh1a2*^-/-^ condition. The different sensitivity of distinct RA response elements (RAREs) to RA levels would enable cells to switch on different gene repertoires depending on RA concentration. The strategy of using binding sites varying in affinity is adopted, for example, by the morphogen Bicoid in fly embryos (Driever et al., 1989). RA itself forms a concentration gradient during development (Shimozono et al., 2013) and such different RAREs could be used to generate positional information by translating RA concentrations into distinct gene expression profiles.

In order to safeguard their developmental capabilities, pluripotent cells, such as the ones comprising the epiblast/PS-like population, should protect themselves from the neuralizing action of RA and therefore need to carefully control the RA levels they are exposed to. This can in theory be achieved at several levels either by controlling the availability of its substrate, the RA synthesis steps, or RA degradation itself (Rhinn and Dollé, 2012). We determined that the Aldh1a2-indipendent RA synthesis could not be attributed to a single aldehyde dehydrogenase by systematically knocking out all the expressed ones (data not shown). Furthermore, we could not identify a combination of knockouts of aldehyde dehydrogenases able to completely abrogate RA synthesis. Tuning RA synthesis can also be achieved by controlling the expression of retinol dehydrogenases that oxidize vitamin A into retinal (Rhinn and Dollé, 2012). Rdh10 could be such a candidate but its expression starts to be tissue-specific only at later developmental stages (Sandell et al., 2007). An additional mechanism adopted by the PS-like cells is the regulation of vitamin A availability through the Rbp1-Stra6 axis. The specific upregulation of the retinol binding protein Rbp1 in the PS-like cells is in accordance with its expression in the mouse embryo (Ruberte et al., 1991; Peng et al., 2019). The control of RA synthesis, however, does not protect the producing cells from the differentiation effects elicited by RA. Our co-culture experiment showed that the response to RA in an RA-producing population was reduced by the presence in the same dish of cells with impaired RA synthesis. Therefore, a cell’s response was not intrinsically determined by its own RA production. The ability to degrade RA via cytocrome P450 Cyp26 enzymes offers an important checkpoint to control the penetrance of a population’s response to RA. Indeed, the knockout of Cyp26a1 resulted in a larger fraction of cells expressing the RA activity reporter. Cyp26a1 mRNA levels were upregulated in the T^TagBFP+^ compared to the Sox1^GFP+^ population, matching the expression in the PS in mouse embryos around E7.0 (Fujii et al., 1997; Peng et al., 2019). Cyp26a1-mediated RA degradation was crucial in limiting the differentiation towards the neural lineage of wild type and *Chrd*^-/-^ *Nog*^-/-^ or *Dkk1*^-/-^ PS-like cells. Altogether, cells exploited a three-tiered control of RA levels regulating precursor availability, RA synthesis and degradation in order to induce neural differentiation only in a subpopulation of cells.

Another crucial aspect highlighted by our study about the control of neural fate acquisition by RA is the role of its receptors (RARs). Confirming previous *in vivo* studies (Mic et al., 2003), the RAR family of RA receptors was responsible for the transduction of RA signaling and importantly to elicit RA induced neural differentiation. This was demonstrated by the absence of expression of the RA reporter and unchanged proportion of Sox1^GFP+^ cells in the *Rara*^-/-^ *Rarb*^-/-^ *Rarg*^-/-^ cultures after inhibiting RARs with AGN or promoting RA synthesis with vitamin A. This kind of response was quite different from the one caused by introducing the same perturbations in the *Aldh1a2*^-/-^ cells, demonstrating that only the RAR-null condition was completely devoid of RA signaling in our system. Notably, the absence of RA signaling due to the deletion of receptors did not prevent the formation of neural progenitors. These results seem at first glance in contradiction with the possibility to reduce neuroectoderm formation inhibiting RA signaling with AGN, even in the presence of Wnt or TGF*β* antagonists. The discrepancies could derive from considering RARs only as ligand dependent transcriptional activators. Instead, RARs can bind to their cognate RAREs in the absence of ligand (Chambon, 1996), repressing target gene expression. We propose that they might act as dams gating the expression of genomic loci important for neural specification and RA presence would lead to a ligand-mediated release of this repression. Their physical absence in RAR-null cells would remove these dams and could additionally unmask binding sites, making them available to other nuclear receptors and transcription factors. In this perspective, both AGN treated cells and RAR-null cells lack the component of positive regulation of the RA targets. However, the *Rara*^-/-^ *Rarb*^-/-^ *Rarg*^-/-^ cells suffer of the additional absence of the ligand-independent receptor repressive function. Two relevant examples of aberrant gene regulation able to promote increased neuroectoderm formation in the RAR-null cells subjected to the PS-like differentiation are the upregulation of Sox2 and downregulation of Tbx6 compared to wild type and AGN-treated cells. Receptor-mediated repression of neural fate acquisition was especially highlighted by the fact that in absence of RA the vast majority of RAR-null cells differentiated to Sox1^GFP+^ neural progenitors, whereas only a fraction of the cells with functional RARs expressed GFP. Altogether, the differentiation of the triple RAR knockout highlighted a more complex involvement of the RA pathway in neuroectoderm differentiation, notwithstanding the intrinsic limitation of removing both activator and repressor functions of the receptors.

In conclusion, we dissected the involvment of RA signaling in the formation of neuroectodermal fate starting from epiblast and PS-like cells. The flexibility of our *in vitro* system allowed us to manipulate the external environment in a controlled manner and interpret classical mouse genetic experiments in a new light. We identified the presence of residual RA signaling in the *Aldh1a2*^-/-^ cells thanks to a sensitive RA reporter. We demonstrated that Cyp26a1 is crucial to safeguard the developmental capabilities of the PS-like cells, even in the *Chrd*^-/-^ *Nog*^-/-^ or *Dkk1*^-/-^ cells. The triple RAR knockout and the possibility to counteract neuroectoderm formation by blocking the RA receptors with AGN, even in combination with TGF*β* or Wnt antagonists, suggest that RA receptors control at least in a negative way key loci for neural induction. Overall, we propose two mechanisms whereby components of the RA pathway can control the formation of neural progenitors: a RA-dependent neural induction gated by RARs and a receptor-mediated repression in the absence of ligand. Our results highlight the potential of studying the self-regulation of fates in absence of spatial organization in ESC-based systems to gain new insights about fundamental aspects of mammalian development.

## Supporting information

Supplementary Information consisting of 6 Supplementary Figures

## Author Contributions

L.R. designed experiments, performed most experiments described in the manuscript and analyzed data. H.L.S. provided critical preliminary data. P.A.N. conceived and supervised the study, performed experiments and analyzed data. L.R. and P.A.N. wrote the paper and H.L.S. commented on the manuscript.

## Acknowledgements

We thank Lucia Cassella for advice on RNA-seq data analysis. We thank Laura Villacorta for help with scRNA-Seq sample processing. This work was technically supported by the EMBL Flow Cytometry Core and Genomics Core facilities. The study was funded by EMBL. L.R. was also supported by the EMBL International PhD Programme (EIPP).

## EXPERIMENTAL PROCEDURES

### mESC maintenance

The parental mESC line was a *Sox1-Brachyury* double knockin-line (Sladitschek and Neveu, 2019). mESCs were maintained in “LIF+serum” as described previously (Sladitschek and Neveu, 2015b). Briefly, cells were cultured at 37°C with 5% CO_2_ on dishes (Nunc) coated with 0.1% gelatin (Sigma). The pluripotency maintaing medium was prepared as follows: DMEM (high glucose, no glutamine, with sodium bicarbonate)(Invitrogen) supplemented with 15% ES-qualified EmbryoMax Fetal Calf Serum (Millipore), 10 ng/ml murine LIF (EMBL Protein Expression and Purification Core Facility), 1x Non-Essential Amino Acids, 2 mM L-glutamine, 1 mM sodium pyruvate, 100 U/ml penicillin and 100 *μ*g/ml streptomycin, 0.1 mM 2-mercaptoethanol (all Invitrogen). Medium was changed daily and cells were passaged every other day with 0.05% Trypsin-EDTA or StemPro Accutase (Invitrogen) at a passaging ratio of 1/3â–1/12.

### Generation of knockout mESC lines

RNA-guided Cas9 nucleases were used to introduce inactivating mutations in the following genes: *Aldh1a2, Chrd, Cyp26a1, Dkk1, Nog, Rara, Rarb, Rarg, Rbp1*, and *Stra6*.

Guide RNA inserts targeting the fourth exon of *Aldh1a2* (with genome target sequence: 5’-AGGGAGTCATCAAAACCCTG), the fifth exon of *Chrd* (5’-GGTCCGAGTTCTTGGCGCGG), the second exon of *Cyp26a1* (5’-GCGCCCATCACCCGCACCGT), the second exon of *Dkk1* (5’-GATCTGTACACCTCCGACGC), the coding sequence of *Nog* (5’-GGAAGTTACAGATGTGGCTG), the fourth exon of *Rara* (5’-GGTGGGCGAGCTCATTGAGA), the fourth exon of *Rarb* (5’-GCGTGGTGTATTTACCCAGC), the fifth exon of *Rarg* (5’-GTGGGACAAGTTCAGCGAGC), the second exon of *Rbp1* (5’-CACTTTTCGGAACTATATCA or 5’-TCCTGCACGATCTCTTTGTC), the fourth exon of *Stra6* (5’-TCCCCAGCCAAGAAATCCAC), were designed and cloned in pX330-U6-Chimeric-BB-CBh-hSpCas9 following Hsu et al. (2013). The resulting pX330 plasmids were transfected in the appropriate mESC lines using Fugene HD (Promega) according to the manufacturer’s protocol. Successfully edited clones were validated by Sanger sequencing of genomic PCR amplicons.

### Reporter constructs

All constructs were assembled using the MXS-chaining strategy (Sladitschek and Neveu, 2015a). A CAG:: H2B-2xiRFP670-bGHpA cassette (combined with a PGK::NeoR-bGHpA cassette) was used as constitutive fluorescent marker for the co-culture experiments.

Transcriptional reporters consisted of binding sites upstream of a minimal CMV promoter driving the expression of NLS-Scarlet-PEST2D or H2B-Scarlet (both relying on the bright red fluorescent protein mScarlet (Bindels et al., 2017)). The plasmid contained a PGK::HygroR-bGHpA cassette to enable selection with hygromycin. Direct repeats (DR) of RAR binding sites spaced by 5 nucleotides (5’-<u>GGTTCA</u>CCGAA<u>AGTTCA</u>) reported in Rossant et al. (1991) were the base of the regulatory region of the DR5-RARE-Scarlet reporter. Three DR5 spaced by 9 and 10 nucleotides were used. The regulatory sequences of the composite DR reporter cDR-RARE-Scarlet consisted of three RAR binding sites (5’-<u>AGGTCA</u>GA<u>AGTTCAAGGTCA</u>) described in Moutier et al. (2012). Three cDRs spaced by 12 nucleotides were used.

Titration of the response to RA of the cDR-RARE-Scarlet reporter line was performed in N2B27 medium supplemented with all-trans retinoic acid. N2B27 medium was prepared from a 1:1 mixture of DMEM/F12 (without HEPES, with L-glutamine) and neurobasal medium with 0.5x B-27 (without vitamin A) and 0.5x N-2 supplements, 0.25 mM L-glutamine, 0.1 mM 2-mercaptoethanol (all Invitrogen), 10 *μ*g/ml BSA fraction V and 10 *μ*g/ml human recombinant insulin (both Sigma). Fluorescence was measured by flow cytometry 24 h after the addition of RA.

### Transgenic mESC lines

We used the following transgenic cell lines in this study.

- Sox1-GFP, *T*-H2B-3xTagBFP mESCs (Sladitschek and Neveu, 2019).
- Sox1-GFP, *T*-H2B-3xTagBFP, Eomes-H2B-mCherry mESCs (Sladitschek and Neveu, 2019).
- Sox1-GFP, *T*-H2B-3xTagBFP DR5-RARE-NLS-Scarlet-PEST2D-bGHpA mESCs.
- Sox1-GFP, *T*-H2B-3xTagBFP cDR-RARE-NLS-Scarlet-PEST2D-bGHpA mESCs.
- Sox1-GFP, *T*-H2B-3xTagBFP cDR-RARE-H2B-Scarlet-bGHpA mESCs.
- Sox1-GFP, *T*-H2B-3xTagBFP *Aldh1a2*^-/-^ mESCs.
- Sox1-GFP, *T*-H2B-3xTagBFP *Aldh1a2*^-/-^ DR5-RARE-NLS-Scarlet-PEST2D-bGHpA CAG::H2B-2xiRFP670-bGHpA mESCs.
- Sox1-GFP, *T*-H2B-3xTagBFP *Aldh1a2*^-/-^ cDR-RARE-H2B-Scarlet-bGHpA mESCs.
- Sox1-GFP, *T*-H2B-3xTagBFP *Dkk1*^-/-^ mESCs.
- Sox1-GFP, *T*-H2B-3xTagBFP *Chrd*^-/-^ *Nog*^-/-^ mESCs.
- Sox1-GFP, *T*-H2B-3xTagBFP *Cyp26a1*^-/-^ DR5-RARE-NLS-Scarlet-PEST2D-bGHpA mESCs.
- Sox1-GFP, *T*-H2B-3xTagBFP *Rbp1*^-/-^ *Stra6*^-/-^ mESCs.
- Sox1-GFP, *T*-H2B-3xTagBFP *Aldh1a2*^-/-^ *Rbp1*^-/-^ *Stra6*^-/-^ cDR-RARE-H2B-Scarlet-bGHpA mESCs.
- Sox1-GFP, *T*-H2B-3xTagBFP *Cyp26a1*^-/-^ *Chrd*^-/-^ *Nog*^-/-^ mESCs.
- Sox1-GFP, *T*-H2B-3xTagBFP *Cyp26a1*^-/-^ *Dkk1*^-/-^ mESCs.
- Sox1-GFP, *T*-H2B-3xTagBFP *Rara*^-/-^ *Rarb*^-/-^ *Rarg*^-/-^ DR5-RARE-NLS-Scarlet-PEST2D-bGHpA mESCs.

### Primitive streak-like differentiation

For differentiation towards a primitive streak-like fate (Sladitschek and Neveu, 2019), mESCs were seeded at a density of 30-50 cells per mm^2^ (unless reported otherwise) onto 0.1% gelatin coated dishes one day prior to the start of the differentiation procedure. The following day, cells were washed with D-PBS and switched to Advanced RPMI 1640 (ThermoFisher) supplemented with 1 *μ*M IDE-1 (Tocris), 0.2% (v/v) ES cell qualified fetal calf serum (Millipore), 2mM L-glutamine (Sigma). 48 hours after the onset of differentiation, medium was replaced every day.

### Differentiation to mesendoderm fates

For Activin- and CHIR-mediated differentiation, mESCs were seeded at a density of 30 cells per mm^2^ onto 0.1% gelatin coated dishes one day prior to the start of the differentiation procedure. The following day, cells were washed with D-PBS and switched to Advanced RPMI 1640 (ThermoFisher) supplemented with 0.2% (v/v) fetal calf serum (Millipore), 2 mM L-glutamine (Sigma) and 3 *μ*M CHIR99021 (Tocris) or 50 ng/ml Activin A (Peprotech). 48 hours after the onset of differentiation, medium was replaced every day.

### Differentiation to neural progenitors for transcriptional profiling

For retinoic acid-mediated differentiation to neural progenitors, mESCs were seeded at a density of 100-200 cells per mm^2^ onto 0.1% gelatin coated dishes one day prior to the start of the differentiation procedure. The following day, cells were washed with D-PBS and switched to N2B27 medium (N2B27 medium was prepared from a 1:1 mixture of DMEM/F12 (without HEPES, with L-glutamine) and neurobasal medium with 0.5x B-27 (with vitamin A) and 0.5x N-2 supplements, 0.25 mM L-glutamine, 0.1 mM 2-mercaptoethanol (all Invitrogen), 10 *μ*g/ml BSA fraction V and 10 *μ*g/ml human recombinant insulin (both Sigma)). *all-trans*-Retinoic acid (Sigma) was added at 1 *μ*M (unless stated otherwise) to the differentiation medium 24 h after the start of the differentiation procedure. Medium was replaced every other day. For the *Cyp26a1*^-/-^ cells and RAR-null cells differentiation in N2B27 medium, B27 supplement without vitamin A was used instead.

For differentiation to neural progenitors mediated by TGF*β* signaling inhibition, mESCs were seeded at a density of 30 cells per mm^2^ onto 0.1% gelatin coated dishes one day prior to the start of the differentiation procedure. The following day, cells were washed with D-PBS and switched to Advanced RPMI 1640 (ThermoFisher) supplemented with 0.2% (v/v) ES cell qualified fetal calf serum (Millipore), 2mM L-glutamine (Sigma) and 10 *μ*M SB431542 (Tocris). 3 *μ*M CHIR99021 (Tocris) was added to the medium from day 3 onwards. 48 hours after the onset of differentiation, medium was replaced every day.

For differentiation to neural progenitors mediated by Wnt signaling inhibition, mESCs were seeded at a density of 30 cells per mm^2^ onto 0.1% gelatin coated dishes one day prior to the start of the differentiation procedure. The following day, cells were washed with D-PBS and switched to Advanced RPMI 1640 (ThermoFisher) supplemented with 0.2% (v/v) ES cell qualified fetal calf serum (Millipore), 2mM L-glutamine (Sigma) and 1 *μ*M IDE-1 and 1 *μ*M XAV939 (both Tocris). 48 hours after the onset of differentiation, medium was replaced every day.

### Pharmacological treatments

For pharmacological interference with PS-like differentiation, compounds were added after 3 days of differentiation (unless indicated otherwise) to Advanced RPMI 1640 (ThermoFisher) supplemented with 1 *μ*M IDE-1 (Tocris), 0.2% (v/v) fetal calf serum (Millipore), 2mM L-glutamine (Sigma). Recombinant Activin A, DKK1, Noggin, sFRP1 were obtained from Peprotech and used at 50 ng/ml from the onset of the PS-like differentiation procedure. All-trans retinoic acid (Sigma) and vitamin A (all-trans retinol, Sigma) were used at 1 *μ*M and 70 nM respectively unless indicated otherwise. AGN193109 (200 nM), CHIR99021 (3 *μ*M), SB431542 (10 *μ*M), XAV939 (1 *μ*M) were all obtained from Tocris and used at the indicated concentrations unless stated otherwise.

### Imaging

Reporter fluorescence was assessed in live differentiating cultures. Confocal images were acquired on an inverted SP8 confocal microscope (Leica) equipped with a 40x PL Apo 1.1 W objective and an incubation chamber at 37C and 5% CO_2_.

### Flow cytometry and fluorescence-activated cell sorting (FACS)

Cells were trypsinized, dissociated to single-cell suspension, pelleted at 1000g for 1 min, resuspended in D-PBS and strained through a 40 *μ*m filter. Samples were analyzed on an LSRFortessa flow cytometer (BD BioSciences). Cells were FACS-purified according to their TagBFP, GFP or Scarlet fluorescence levels using an Aria Fusion sorter (BD BioSciences). Flow cytometry data was analyzed with FlowJo.

### RNA-seq library construction

RNA was extracted from pellets of trypsinized cells using the MirVana kit (Ambion) following the instructions provided by the manufacturer. Barcoded mRNA libraries were prepared using TruSeq RNA Sample Preparation (Illumina) following the manufacturer’s instructions. The libraries were sequenced on Illumina NextSeq 500 in the high density 75 bp single-end regime. Sequencing results are available on ArrayExpress with accession number E-MTAB-10242. In addition, we used mRNA expression data that we previously deposited on ArrayExpress with accession E-MTAB-2830, E-MTAB-3234 and E-MTAB-4904.

### RNA-seq analysis

Ensembl cDNAs of the mouse genome release GRCm38 were masked with RepeatMasker (Smit, AFA, Hubley, R and Green, P. RepeatMasker Open-3.0. 1996-2010 http://www.repeatmasker.org) and a Bowtie index was built using these masked transcripts. Reads were aligned to this index using Bowtie (Langmead et al., 2009) with default parameters. mRNA read counts were determined for each Ensembl ID by parsing the Bowtie output.

The differential gene expression analysis was performed using the Bioconductor package edgeR (Robinson et al., 2010). Starting from raw read counts, a normalization factor was applied taking into account differences in sequencing depth and effective library size among the libraries. Providing experimental design matrix, dispersion estimates were obtained and negative binomial generalized linear models (GLMs) were fitted to the read counts. A quasi-likelihood (QL) F-test was then applied to determine differential expression (DE) across the conditions. The KEGG pathway enrichment analysis was performed to identify over- or under-represented pathways, starting from the list of DE genes, using the edgeR built-in function kegga.

To identify transcription regulators differentially expressed in the four neural progenitors differentiation procedures, transcription regulators with FDR<0.05, maximal read counts >4 RPM and fold change >8 between the assessed samples were selected. Genes with differential expression between cDR-RARE-Scarlet positive and negative cells were selected with the criteria FDR<0.05, maximal read counts >4 RPM and fold change >2 between the assessed samples. To identify transcription regulators differentially expressed in *Rara*^-/-^ *Rarb*^-/-^ *Rarg*^-/-^ cells, we used as criteria FDR<0.05, maximal read counts >4 RPM and fold change >2 between the assessed samples.

Principal component analysis was carried out as described in Neveu et al. (2010).

### Single-cell RNA-Sequencing

Cultures undergoing PS-like differentiation were trypsinized at day 2 to 5. 8 different samples were collected: entire cultures at day 2, 3, 4 and 5 (day 5 was represented in biological triplicates) and FACS-purified TagBFP+/GFP- and GFP+ subpopulations from day 4. The solution was pelleted for 1 min at 1,000g and resuspended in D-PBS+0.04% Bovine Serum Albumin and strained through a 40 *μ*m filter. For each sample, 8,000 to 10,000 cells of a single-cell suspension (of concentration 1,000 cells/*μ*l) were loaded on a Chromium Controller (10x Genomics). Libraries were prepared using the Chromium Single Cell 3’ Reagent Kit (10x Genomics) with v3 Chemistry according to the manufacturer’s instructions. The eight barcoded libraries were sequenced in four runs on Illumina NextSeq 500 in the high density 40 bp paired-end regime. scRNA-Seq results are available on ArrayExpress with accession number E-MTAB-10243.

### scRNA-seq analysis and quality control

For each sample, reads were demultiplexed according to the cell barcodes. mRNA reads were aligned to a mouse cDNA index using Bowtie (Langmead et al., 2009) allowing up to 3 mismatches. The Bowtie output was parsed to count the number of unique molecular identifiers (UMIs) and reads aligning to each transcript model. We kept cells with >5,000 UMIs, >2,000 expressed genes and a fraction of mitochondrial transcript <8%. For each library, cells with UMI counts greater than the average UMI count plus three standard deviations were discarded in order to remove doublets. Overall, 46,700 cells in total for the eight libraries passed quality controls. Expression levels for individual cells were normalized using Seurat methods (Satija et al., 2015). Dimensionality reduction was performed using uniform manifold approximation and projection (UMAP) (Becht et al., 2018). Clustering into subpopulations was performed using Ward distance after dimensionality reduction by UMAP. To identify potential markers of the different cell categories, we retained genes with at least 25% expressing cells and an average expression of 8 RPM in one cell category.

### Statistical analysis

Statistical tests were computed using R or the Python SciPy module.

### Data availability

Sequencing results are deposited on ArrayExpress with accession numbers E-MTAB-10242 and E-MTAB-10243. In addition, we used mRNA expression data that we previously deposited on ArrayExpress with accession E-MTAB-2830, E-MTAB-3234 (Sladitschek and Neveu, 2015b) and E-MTAB-4904 (Sladitschek and Neveu, 2019).

### Supplemental Information

Supplemental Information includes six figures.

