## Supplementary Information consisting of 6 Supplementary Figures for "Multi-layered regulation of neuroectoderm differentiation by retinoic acid in a primitive streak-like context"

#### Figure S1

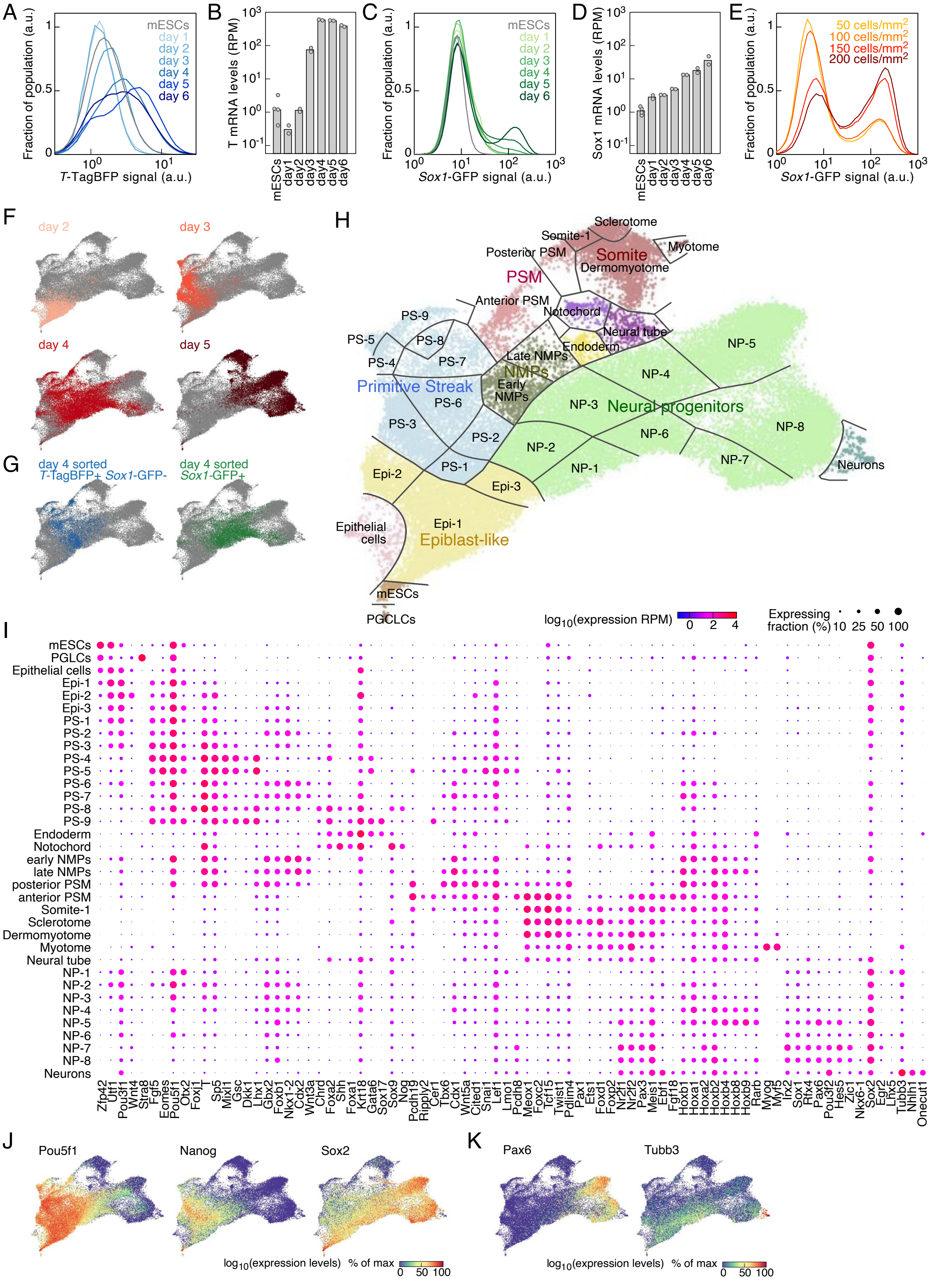

A
Fraction of population (a.u.)
T-TagBFP signal (a.u.)
mESCs
day 1
day 2
day 3
day 4
day 5
day 6
B
T mRNA levels (RPM)
mESCs
day1
day2
day3
day4
day5
day6
C
Fraction of population (a.u.)
Sox1-GFP signal (a.u.)
mESCs
day 1
day 2
day 3
day 4
day 5
day 6
D
Sox1 mRNA levels (RPM)
mESCs
day1
day2
day3
day4
day5
day6
E
Fraction of population (a.u.)
Sox1-GFP signal (a.u.)
50 cells/mm2
100 cells/mm2
150 cells/mm2
200 cells/mm2
F
day 2
day 3
day 4
day 5
G
day 4 sorted T-TagBFP+ Sox1-GFP-
day 4 sorted Sox1-GFP+
H
Sclerotome
Somite-1
Myotome
Posterior PSM
Somite
Dermomyotome
PSM
Anterior PSM
Notochord
Neural tube
Endoderm
Late NMPs
NMPs
Early NMPs
PS-9
PS-8
PS-7
PS-6
PS-5
PS-4
PS-3
PS-2
PS-1
Primitive Streak
NP-1
NP-2
NP-3
NP-4
NP-5
NP-6
NP-7
NP-8
Neural progenitors
Neurons
Epi-1
Epi-2
Epi-3
Epiblast-like
Epithelial cells
mESCs
PGCLCs
I
log10(expression RPM)
Expressing fraction (%) 10 25 50 100
mESCs
PGLCs
Epithelial cells
Epi-1
Epi-2
Epi-3
PS-1
PS-2
PS-3
PS-4
PS-5
PS-6
PS-7
PS-8
PS-9
Endoderm
Notochord
early NMPs
late NMPs
posterior PSM
anterior PSM
Somite-1
Sclerotome
Dermomyotome
Myotome
Neural tube
NP-1
NP-2
NP-3
NP-4
NP-5
NP-6
NP-7
NP-8
Neurons
Zfp42 Utf1 Pou3f1 Wnt4 Stra8 Fgf5 Eomes Pou5f1 Otx2 Foxj1 T Sp5 Mixl1 Gsc Dkk1 Lhx1 Gbx2 Foxb1 Nkx1-2 Cdx2 Wnt3a Chrd Foxa2 Shh Foxa1 Krt18 Gata6 Sox17 Sox9 Nog Pcdh19 Ripply2 Cer1 Tbx6 Cdx1 Wnt5a Cited1 Snai1 Lef1 Lmo1 Pcdh8 Meox1 Foxc2 Tcf15 Twist1 Pdlim4 Pax1 Ets1 Foxd1 Foxp2 Nr2f1 Nr2f2 Pax3 Meis1 Ebf1 Fgf18 Hoxb1 Hoxa1 Hoxa2 Hoxb2 Hoxb4 Hoxb8 Hoxb9 Rarb Myog Myf5 Irx2 Sox1 Rfx4 Pax6 Pou3f2 Hes5 Zic1 Nkx6-1 Sox2 Egr2 Lhx5 Tubb3 Nhlh1 Onecut1
J
Pou5f1
Nanog
Sox2
log10(expression levels) % of max
0 50 100
K
Pax6
Tubb3
log10(expression levels) % of max
0 50 100

**Figure S1. Related to Figure 1. Characterization of neural induction by primitive streak-like cells at single cell level.**
(A) Time course of *T*-TagBFP expression levels in cultures undergoing PS-like differentiation (gray: mESCs, blue line color according to the day of differentiation).
(B) T mRNA levels in cultures undergoing PS-like differentiation (RPM: reads per million mapped reads).
(C) Time course of *Sox1*-GFP expression levels in cultures undergoing PS-like differentiation (gray: mESCs, green line color according to the day of differentiation).
(D) Sox1 mRNA levels in cultures undergoing PS-like differentiation (RPM: reads per million mapped reads).
(E) *Sox1*-GFP expression levels in cultures undergoing PS-like differentiation with increasing starting cell density (yellow: 50 cells/mm$^2$, orange: 100 cells/mm$^2$, red: 150 cells/mm$^2$, dark red: 200 cells/mm$^2$).
(F) UMAP colored by the day of differentiation.
(G) UMAP location of cells FACS-purified according to their *T*-TagBFP and *Sox1*-GFP expression at day 4.
(H) UMAP colored according to the identified populations (NMPs: neuromesodermal progenitors, NP: neural progenitors, PSM: presomitic mesoderm, PGCLCs: primordial germ cell-like cells, PS: Primitive streak).
(I) Dot plots of marker expression levels in the 35 identified populations (RPM: reads per million mapped reads).
(J) UMAP colored by the scaled expression of pluripotency markers.
(K) UMAP colored by the scaled expression of the neural progenitor marker Pax6 and the neuronal marker Tubb3.

### Figure S2

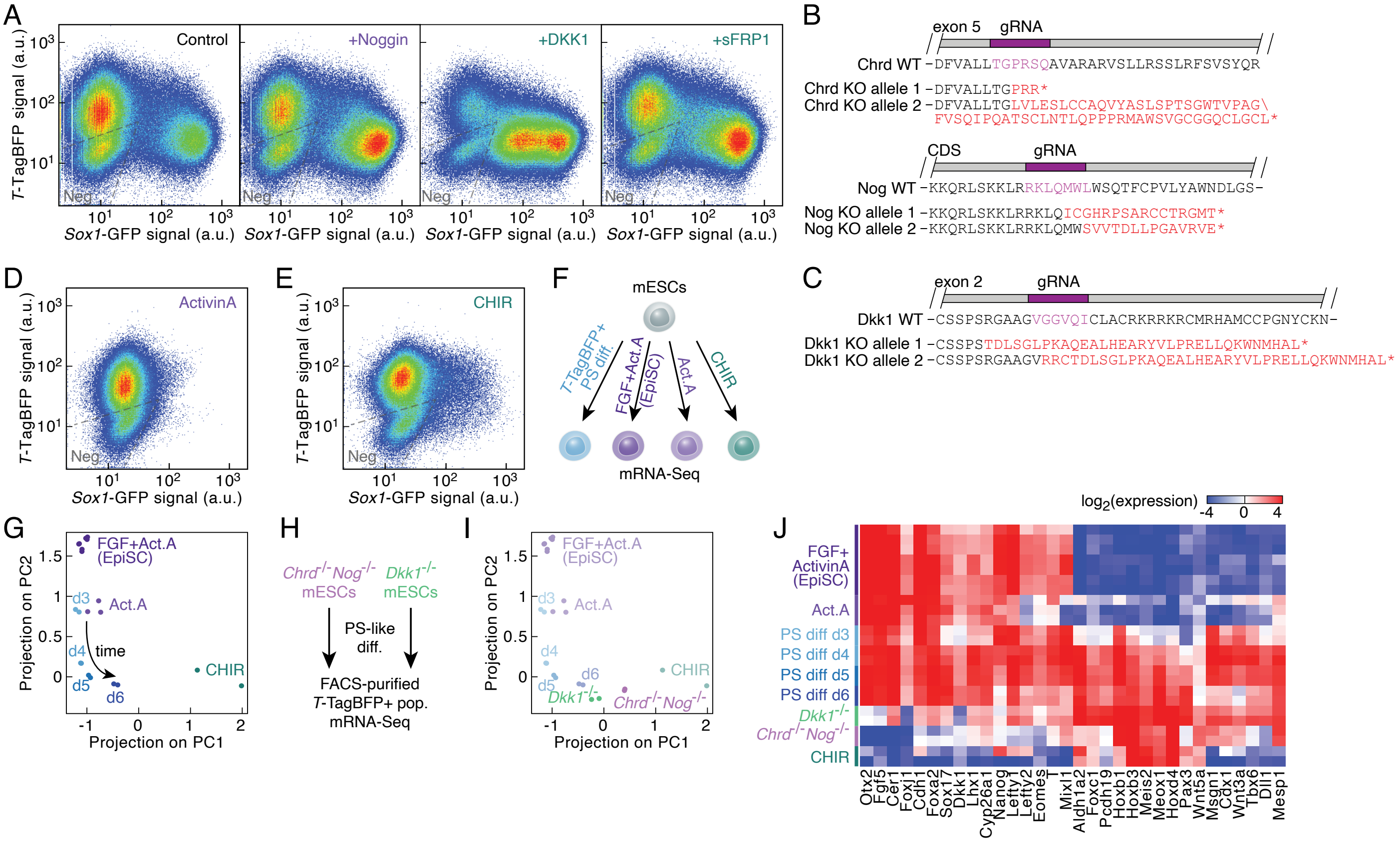


**Figure S2. Related to Figure 2. A balance between agonists and inhibitors of the TGF$\beta$ and Wnt signaling pathways fine tunes the formation of neuroectodermal and primitive streak derivatives in culture.**

(A) *T*-TagBFP and *Sox1*-GFP reporter expression after 5 days of PS-like differentiation (left panel) supplemented with recombinant Noggin (middle left panel), DKK1 (middle right panel) or sFRP1 (right panel).

(B) Sanger sequencing-validated obtained alleles of *Chrd*$^{-/-}$ *Nog*$^{-/-}$ mESCs. The relative position of the guide RNA used to target the locus is indicated in purple. $\star$: stop codon.

(C) Sanger sequencing-validated obtained alleles in *Dkk1*$^{-/-}$ mESCs. The relative position of the guide RNA used to target the locus is indicated in purple. $\star$: stop codon.

(D, E) *T*-TagBFP and *Sox1*-GFP reporter expression in cultures differentiated for 4 days with Activin A (E) or CHIR (F).

(F) Experimental strategy to characterize the impact of the activation of the TGF$\beta$ and Wnt signaling pathways on PS-like populations. Epiblast Stem Cells (EpiSCs, maintained with FGF and Activin A) serve as an additional comparison point.

(G) Principal component analysis of expression profiles of PS-induced T$^{\text{TagBFP+}}$ cells (blue, color according to the day of differentiation), cells after Activin A (purple) or CHIR (teal blue) differentiation and EpiSCs (dark purple).

(H) Scheme to characterize the fate induced by the PS-like differentiation of *Chrd*$^{-/-}$ *Nog*$^{-/-}$ or *Dkk1*$^{-/-}$ mESCs.

(I) Projection on the first two principal components of expression profiles of T$^{\text{TagBFP+}}$ *Chrd*$^{-/-}$ *Nog*$^{-/-}$ (pink) and *Dkk1*$^{-/-}$ (light green) cells after PS-like differentiation.

(J) Expression levels of primitive streak, endoderm, presomitic mesoderm and somite markers in EpiSCs (dark purple), cells differentiated with Activin A (purple) or CHIR (teal blue), $T^{TagBFP+}$ wild type cells (PS diff d3, d4, d5 and d6) after PS-like differentiation, $T^{TagBFP+}$ *Chrd*$^{-/-}$ *Nog*$^{-/-}$ (pink) and *Dkk1*$^{-/-}$ (light green) cells after PS-like differentiation.

#### Figure S3

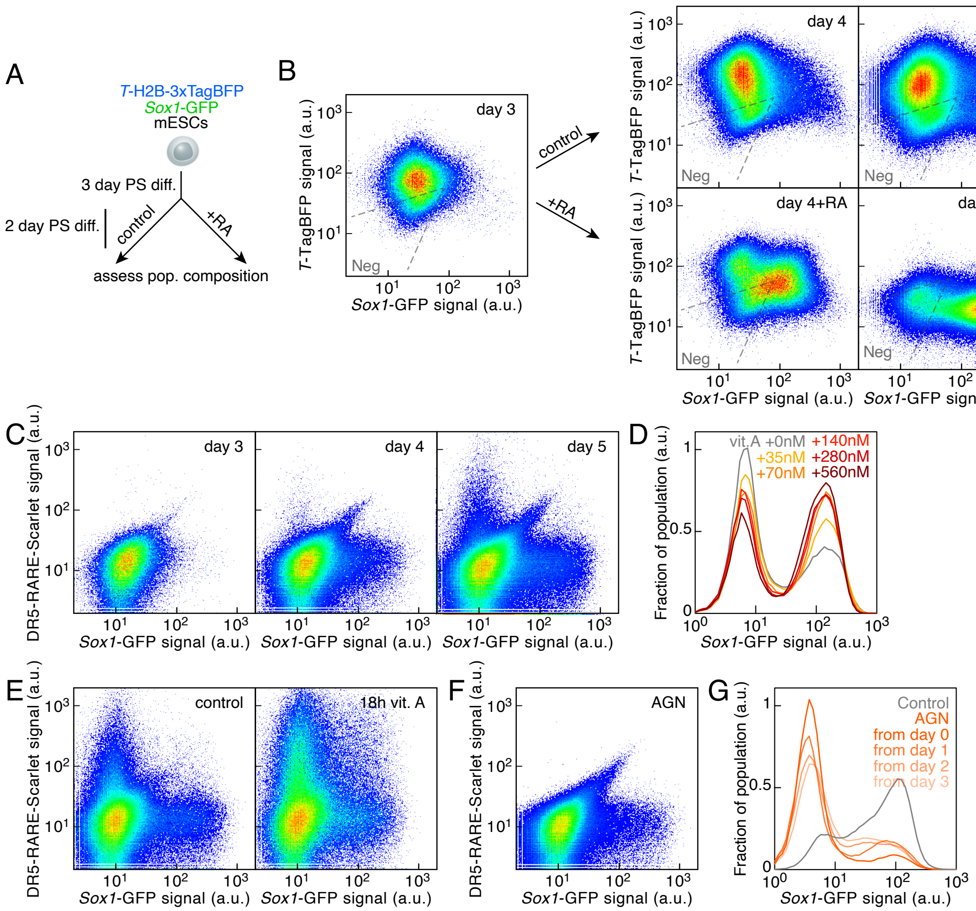


**Figure S3. Related to Figure 3. Retinoic acid signaling contributes to the formation of neural progenitors during the by PS-like differentiation.**

(A) Experimental strategy to monitor the impact of exogenous RA on the PS-like differentiation.

(B) Time course of *T*-TagBFP and *Sox1*-GFP expression levels without (top row) or with supplementation of 1 $\mu$M RA (bottom row) after day 3 of PS-like differentiation (left panel, middle panels: day 4, right panel: day 5).

(C) Time course of the expression levels of *Sox1*-GFP and a DR5-based RA signaling reporter (DR5-RARE::Scarlet) in cultures undergoing PS-like differentiation (left: day 3, middle: day 4, right: day 5).

(D) *Sox1*-GFP expression levels in cultures undergoing PS-like differentiation in medium with additional vitamin A (vit. A), a precursor of RA (gray: no additional vit. A, line color according to the indicated concentration in nM).

(E) *Sox1*-GFP and DR5-RARE-Scarlet expression levels after PS-like differentiation (left: control, right: addition of 35 nM vitamin A for 18 h).

(F) *Sox1*-GFP and DR5-RARE-Scarlet expression levels after 5 days of PS-like differentiation with the addition of the RAR antagonist AGN for the last 48 hours before the analysis.

(G) *Sox1*-GFP expression levels in cultures undergoing PS-like differentiation in presence of the RAR antagonist AGN (gray: control, orange line color according to starting day of AGN treatment).

#### Figure S4

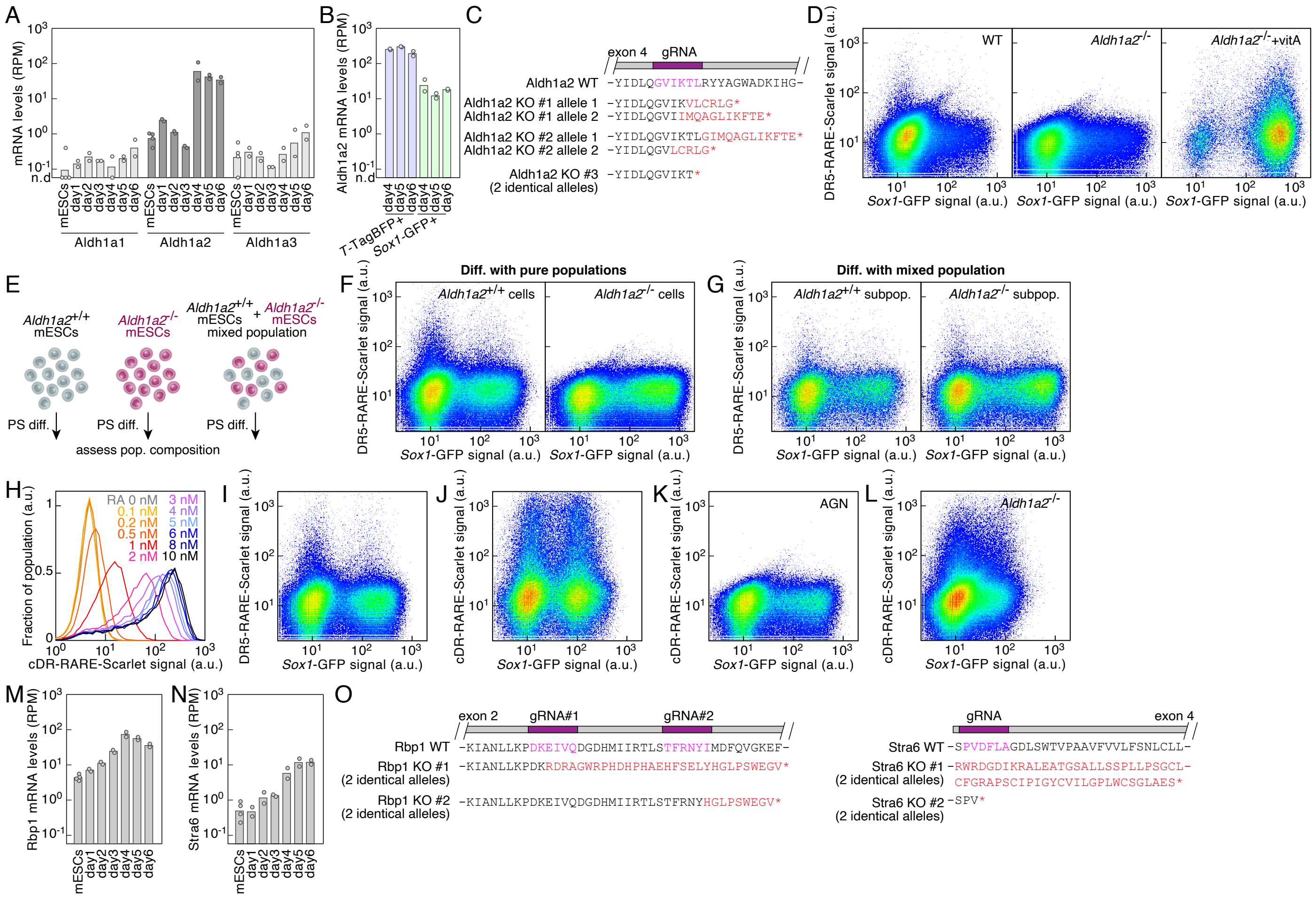


**Figure S4. Related to Figure 4. Aldh1a2-independent RA signaling during PS-like differentiation.**
(A) mRNA expression time course of Aldh1a1, Aldh1a2, Aldh1a3 during PS-like differentiation (RPM: reads per million mapped reads).
(B) Aldh1a2 mRNA expression levels in $T^{TagBFP+}$ and $Sox1^{GFP+}$ cells during PS-like differentiation (RPM: reads per million mapped reads).
(C) Sanger sequencing-validated obtained alleles of *Aldh1a2*$^{-/-}$ cells. The relative position of the guide RNA used to target the locus is indicated in purple. ⋆: stop codon.
(D) *Sox1*-GFP and DR5-RARE-Scarlet reporter expression after PS-like differentiation of wild type (left panel) or *Aldh1a2*$^{-/-}$ cells without (middle panel) or with additional vitamin A (right panel).
(E) Scheme of the experimental principle to assess whether RA signaling acts in a cell-autonomous manner during the PS-like differentiation by mixing wild type and *Aldh1a2*$^{-/-}$ mESCs that can be distinguished by the expression of a constitutive fluorescent marker.
(F, G) *Sox1*-GFP and DR5-RARE-Scarlet reporter expression after PS-like differentiation of pure wild type (F, left panel) or *Aldh1a2*$^{-/-}$ (F, right panel) cultures or a mixed population (G) containing wild type (G, left panel) and *Aldh1a2*$^{-/-}$ (G, right panel) cells.
(H) cDR-RARE-Scarlet reporter expression after 24 hour treatment with RA (line color according to the indicated concentrations -between 0 and 10 nM).
(I) *Sox1*-GFP and DR5-RARE-Scarlet reporter expression after PS-like differentiation of wild type cells.
(J) *Sox1*-GFP and cDR-RARE-Scarlet reporter expression after PS-like differentiation of wild type cells.

(K) *Sox1*-GFP and cDR-RARE-Scarlet reporter expression after PS-like differentiation of wild type cells with the RAR antagonist AGN.
(L) *Sox1*-GFP and cDR-RARE-Scarlet reporter expression after PS-like differentiation of *Aldh1a2*$^{-/-}$ cells.
(M) Rbp1 mRNA expression time course during PS-like differentiation (RPM: reads per million mapped reads).
(N) Stra6 mRNA expression time course during PS-like differentiation (RPM: reads per million mapped reads).
(O) Sanger sequencing-validated obtained alleles of *Rbp1*$^{-/-}$*Stra6*$^{-/-}$ mESCs (clone ♯1 in wild type cells, clone ♯2 in *Aldh1a2*$^{-/-}$ cells). The relative position of the guide RNA used to target the locus is indicated in purple. ⋆: stop codon.

#### Figure S5

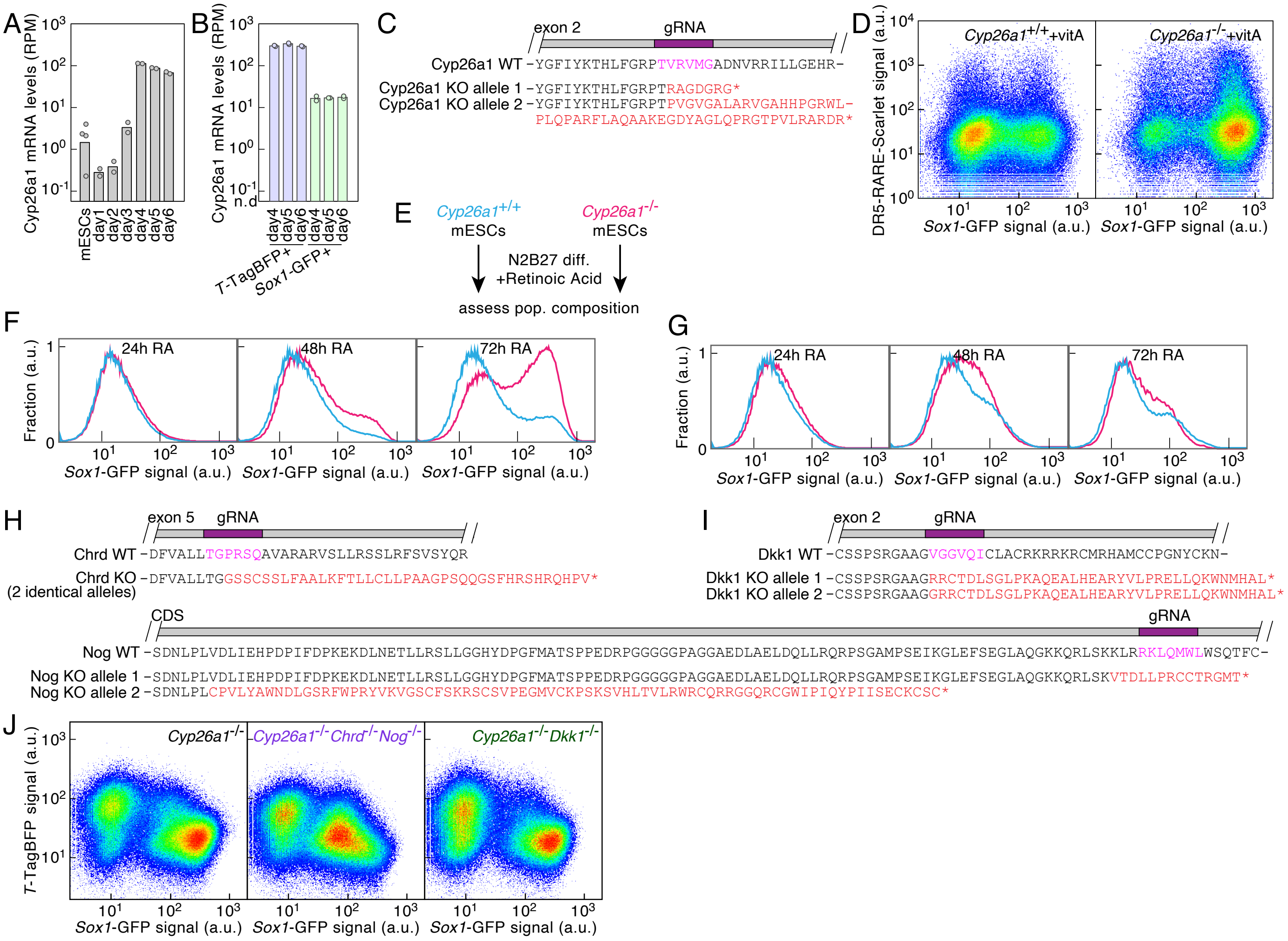


**Figure S5. Related to Figure 5. Cyp26a1 is a key factor limiting RA levels and neuroectoderm differentiation during PS-like differentiation.**
(A) Cyp26a1 mRNA levels in cultures undergoing PS-like differentiation (RPM: reads per million mapped reads).
(B) Cyp26a1 mRNA expression levels in $T^{TagBFP+}$ and $Sox1^{GFP+}$ cells during PS-like differentiation (RPM: reads per million mapped reads).
(C) Sanger sequencing-validated obtained alleles of *Cyp26a1*$^{-/-}$ mESCs. The relative position of the guide RNA used to target the locus is indicated in purple. ⋆: stop codon.
(D) *Sox1*-GFP and DR5-RARE-Scarlet reporter expression after PS-like differentiation of wild type (left panel) or *Cyp26a1*$^{-/-}$ cells (right panel) with additional vitamin A from day 3.
(E) Scheme of the experimental principle to assess the impact of Cyp26a1 knockout on the course of neuroectoderm differentiation in chemically defined medium (N2B27).
(F) *Sox1*-GFP reporter expression after differentiation with 1 nM RA for 24, 48 and 72 hours of wild type cells (cyan) and *Cyp26a1*$^{-/-}$ (magenta) cells.
(G) *Sox1*-GFP reporter expression after differentiation with 100 nM RA for 24, 48 and 72 hours of wild type cells (cyan) and *Cyp26a1*$^{-/-}$ (magenta) cells.

(H) Sanger sequencing-validated obtained alleles of *Chrd*$^{-/-}$ *Nog*$^{-/-}$ mESCs in a *Cyp26a1*$^{-/-}$ background. The relative position of the guide RNA used to target the locus is indicated in purple. ⋆: stop codon.
(I) Sanger sequencing-validated obtained alleles of *Dkk1*$^{-/-}$ mESCs in a *Cyp26a1*$^{-/-}$ background. The relative position of the guide RNA used to target the locus is indicated in purple. ⋆: stop codon.
(J) *T*-TagBFP and *Sox1*-GFP reporter expression after PS-like differentiation of *Cyp26a1*$^{-/-}$ (left), *Cyp26a1*$^{-/-}$ *Chrd*$^{-/-}$ *Nog*$^{-/-}$ (middle) and *Cyp26a1*$^{-/-}$ *Dkk1*$^{-/-}$ (right) mESCs.

**Figure S6**

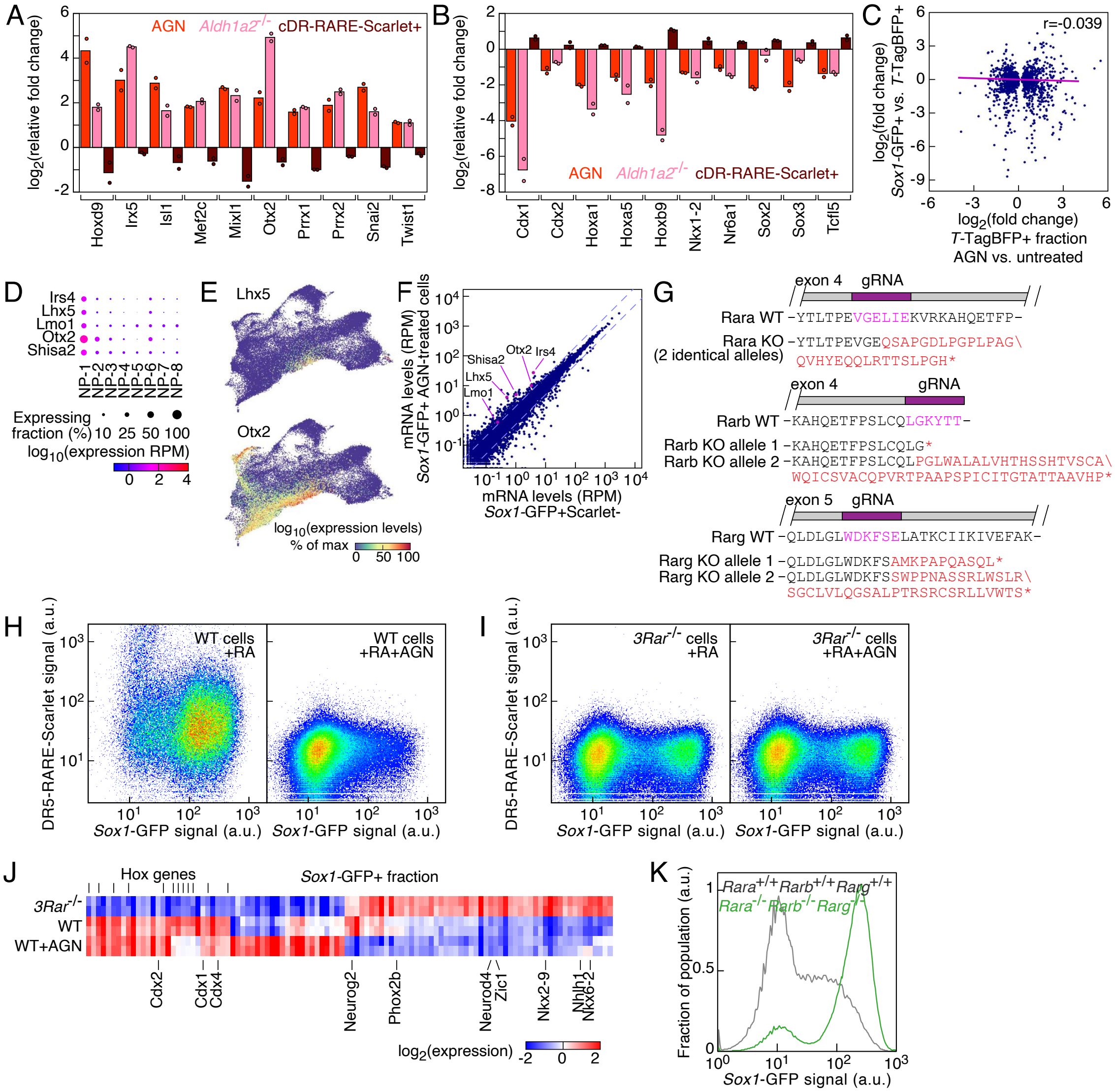


**Figure S6. Related to Figure 6. Role of RA signaling and RAR receptors in neural commitment and in establishing neural progenitor diversity.**
(A, B) Relative changes of expression of transcriptional factors in the $T^{TagBFP+}$ subpopulation of AGN-treated cells, *Aldh1a2*$^{-/-}$ cells or cDR-RARE-Scarlet+ cells compared to untreated wild type $T^{TagBFP+}$ cells (A: genes upregulated by RA signaling block, B: gene downregulated by RA signaling block).
(C) Comparison of differential expression in $T^{TagBFP+}$ cells with or without AGN treatment and the $T^{TagBFP+}$ and $Sox1^{GFP+}$ subpopulations (Pearson's r=-0.039, p=0.14).
(D) Dot plots of markers upregulated in the NP-1 category compared to the other neural progenitor categories (RPM: reads per million mapped reads).
(E) UMAP colored by the scaled expression of Lhx5 or Otx2.
(F) Differential gene expression between $Sox1^{GFP+}$ cells that are cDR-RARE-Scarlet- and AGN-treated $Sox1^{GFP+}$ cells. Dashed lines indicate 2-fold expression changes (RPM: reads per million mapped reads).
(G) Sanger sequencing-validated obtained alleles of *Rara*$^{-/-}$ *Rarb*$^{-/-}$ *Rarg*$^{-/-}$ mESCs. The relative position of

the guide RNA used to target the locus is indicated in purple. ⋆: stop codon.
(H) *Sox1*-GFP and DR5-RARE-Scarlet reporter expression after 4-day differentiation of wild type mESCs to neural progenitors with 100 nM RA (left panel, right panel: 100 nM RA + 1 $\mu$M AGN).
(I) *Sox1*-GFP and DR5-RARE-Scarlet reporter expression after 4-day differentiation of $Rara^{-/-} Rarb^{-/-} Rarg^{-/-}$ mESCs to neural progenitors with 100 nM RA (left panel, right panel: 100 nM RA + 1 $\mu$M AGN).
(J) Expression levels of transcriptional factors with differential expression in the $Sox1^{GFP+}$ fraction after PS-like differentiation of $Rara^{-/-} Rarb^{-/-} Rarg^{-/-}$ ($3Rar^{-/-}$) cells or wild type cells without (WT) or with (WT+AGN) treatment with the RAR antagonist AGN.
(K) *Sox1*-GFP reporter expression after 6 days of differentiation in chemically defined medium (N2B27, without vitamin A) of wild type cells (gray) or $Rara^{-/-} Rarb^{-/-} Rarg^{-/-}$ cells (green).
